# FtsK is Critical for the Assembly of the Unique Divisome Complex of the FtsZ-less *Chlamydia trachomatis*

**DOI:** 10.1101/2024.10.24.620021

**Authors:** McKenna Harpring, Junghoon Lee, Guangming Zhong, Scot P. Ouellette, John V. Cox

**Author notes:** **Correspondence:** Corresponding Author: John V. Cox.

## Abstract

*Chlamydia trachomatis* serovar L2 *(Ct),* an obligate intracellular bacterium that does not encode FtsZ, divides by a polarized budding process. In the absence of FtsZ, we show that FtsK, a chromosomal translocase, is critical for divisome assembly in *Ct*. Chlamydial FtsK forms discrete foci at the septum and at the base of the progenitor mother cell, and our data indicate that FtsK foci at the base of the mother cell mark the location of nascent divisome complexes that form at the site where a daughter cell will emerge in the next round of division. The divisome in *Ct* has a hybrid composition, containing elements of the divisome and elongasome from other bacteria, and FtsK is recruited to nascent divisomes prior to the other chlamydial divisome proteins assayed, including the PBP2 and PBP3 transpeptidases, and MreB and MreC. Knocking down FtsK prevents divisome assembly in *Ct* and inhibits cell division and septal peptidoglycan synthesis. We further show that MreB does not function like FtsZ and serve as a scaffold for the assembly of the *Ct* divisome. Rather, MreB is one of the last proteins recruited to the chlamydial divisome, and it is necessary for the formation of septal peptidoglycan rings. Our studies illustrate the critical role of chlamydial FtsK in coordinating divisome assembly and peptidoglycan synthesis in this obligate intracellular bacterial pathogen.

## Introduction

Most bacteria divide by a highly conserved process termed binary fission, which occurs through the symmetric division of the parental cell into two daughter cells (Harpring 2023). However, *Chlamydia trachomatis* serovar L2 (*Ct*), a coccoid, gram-negative, obligate intracellular bacterium divides by a polarized cell division process characterized by an asymmetric expansion of the membrane from one pole of a coccoid cell resulting in the formation of a nascent daughter cell (Abdelrahman 2016; Ouellette SP 2022).

*Ct* undergoes a biphasic developmental cycle during infection. Non-replicating and infectious elementary bodies (EBs) bind to and are internalized by target cells. Following internalization, EBs within a membrane vacuole, termed the inclusion, differentiate into replicating reticulate bodies (RBs). After replication, RBs undergo secondary differentiation into EBs, which are released from cells to initiate another round of infection (Abdelrahman 2005).

In evolving to obligate intracellular dependence, *Ct* has eliminated several gene products essential for cell division in other bacteria, including the central coordinator of divisome formation, FtsZ (Stephens 1998; Ouellette SP 2020). This tubulin-like protein forms filaments that associate to form a ring at the division plane (Barrows 2021), which serves as a scaffold for the assembly of the other components of the bacterial divisome that regulate the processes of septal peptidoglycan (PG) synthesis and chromosomal translocation. Of the twelve divisome proteins shown to be essential for cell division in the model gammaproteobacterial organism, *E. coli, Ct* encodes homologues of FtsK, a chromosomal translocase (Ouellette SP 2012); FtsQLB, regulators of septal PG synthesis (Ouellette SP 2015; Kaur 2022), FtsW, a septal transglycosylase (Putman T 2019), and penicillin binding protein 3 (PBP3/FtsI), a septal transpeptidase (Ouellette SP 2012).

In addition to the divisome, rod-shaped bacteria employ another multiprotein complex, the elongasome, which directs sidewall PG synthesis (Liu 2020) necessary for cell lengthening and the maintenance of cell shape prior to division. Although *Ct* is a coccoid organism, it encodes several elongasome proteins, including MreB, MreC, RodA, RodZ, and penicillin binding protein 2 (PBP2), a sidewall transpeptidase (Ouellette SP 2012; Ouellette SP 2014; Cox 2020). The actin-like protein MreB is essential for cell division (Ouellette SP 2012; Abdelrahman 2016) and forms septal rings in *Ct* (Kemege 2015; Liechti 2016; Lee 2020). These observations led to the proposal that MreB replaces FtsZ in *Ct* and serves as a scaffold necessary for the assembly of the chlamydial divisome (Lee 2020).

While inhibitor studies suggest that chlamydial cell division is dependent upon elements of the divisome and elongasome from other organisms (Ouellette SP 2012; Abdelrahman 2016; Cox 2020), the composition and ordered assembly of the chlamydial divisome and its distribution during polarized budding are undefined. We hypothesized that FtsK, a chromosomal translocase, serves a critical function in regulating the division process of *Ct,* given previous observations demonstrating it interacts with elements of both the elongasome and divisome (Ouellette SP 2012). We show here that FtsK is critical for the assembly of the hybrid divisome complex of *Ct* and that MreB does not serve as a scaffold necessary for the assembly of the chlamydial divisome. Rather, chlamydial MreB associates with this hybrid divisome complex late in the chlamydial divisome assembly process, and MreB filament formation is necessary for the formation of septal PG rings. Therefore, our data identify FtsK as a key regulator of the cell division process of *Ct*.

## Results

### FtsK Forms Foci in *Ct* that Mark the Location of Divisome Complexes

In the *E. coli* linear divisome assembly pathway (Du 2017), FtsK is the first protein downstream of FtsZ encoded by *Ct* (Figure 1A). In other organisms, FtsK is uniformly distributed at the septum of dividing cells (Yu 1998; Wang 2006; Veiga 2017). To investigate the localization of FtsK during cell division in *Ct*, HeLa cells were infected with *Ct*. To overcome the challenges associated with assessing cell morphologies in densely packed inclusions in infected cells, we analyzed FtsK localization in *Ct* derived from lysates of infected HeLa cells at 21 hrs post-infection (hpi) as described previously (Ouellette SP 2022). *Ct* were stained with an antibody against the chlamydial major outer membrane protein (MOMP) and an antibody that recognizes endogenous FtsK. Blotting analysis revealed that this FtsK antibody recognizes a single protein with the predicted molecular mass of FtsK (Supp. Fig. S1A). Our results showed that, unlike FtsK in other organisms, chlamydial FtsK accumulates in discrete foci in the membrane of coccoid cells (Fig. 1B). In cell division intermediates, FtsK localized in foci at the septum, foci at the septum and at the base of the progenitor mother cell, or foci at the base of the progenitor mother cell only (Figure 1C). The chlamydial FtsK foci observed during cell division were not uniformly distributed at the septum, rather septal foci of FtsK were restricted to one side of the MOMP-stained septum. In addition, the FtsK foci were often above or below (marked with arrowheads in Fig. 1C) the MOMP-stained septum. Similar analyses were performed using *Ct* transformed with the pBOMB4-Tet (-GFP) plasmid encoding FtsK with a C-terminal mCherry tag. The expression of this mCherry fusion is under the control of an anhydrotetracycline (aTc)-inducible promoter. HeLa cells were infected with the transformant, and the expression of the fusion was induced by the addition of 10nM aTc to the media of infected cells at 19hpi. RBs were harvested from the induced cells at 21hpi and stained with MOMP antibodies. Imaging analyses revealed that like endogenous FtsK, FtsK-mCherry accumulated in foci in coccoid cells (Fig. 1D), and in division intermediates, it localized in foci at the septum, foci at the septum and at the base of the progenitor mother cell, or foci at the base of the progenitor mother cell only (Fig. 1E). The foci of FtsK-mCherry, like endogenous FtsK, were often offset relative to the plane defined by MOMP staining at the septum (arrowhead in Fig. 1E). Inclusion forming unit (IFU) assays demonstrated that overexpression of the FtsK-mCherry fusion had no effect on chlamydial developmental cycle progression and the production of infectious EBs (Supp. Fig. S2A). While it is possible that the population of FtsK at the base of the mother cell is a remnant of FtsK from a previous division, ∼20% of dividing cells have a secondary bud (Fig. 1F), and FtsK and FtsK-mCherry accumulate in foci at the base of secondary buds (arrowheads in Fig. 1G), suggesting that the population of FtsK at the base of the mother cell corresponds to a nascent divisome complex that forms at the site where the daughter cell will arise in the next round of division.

**Figure 1:**
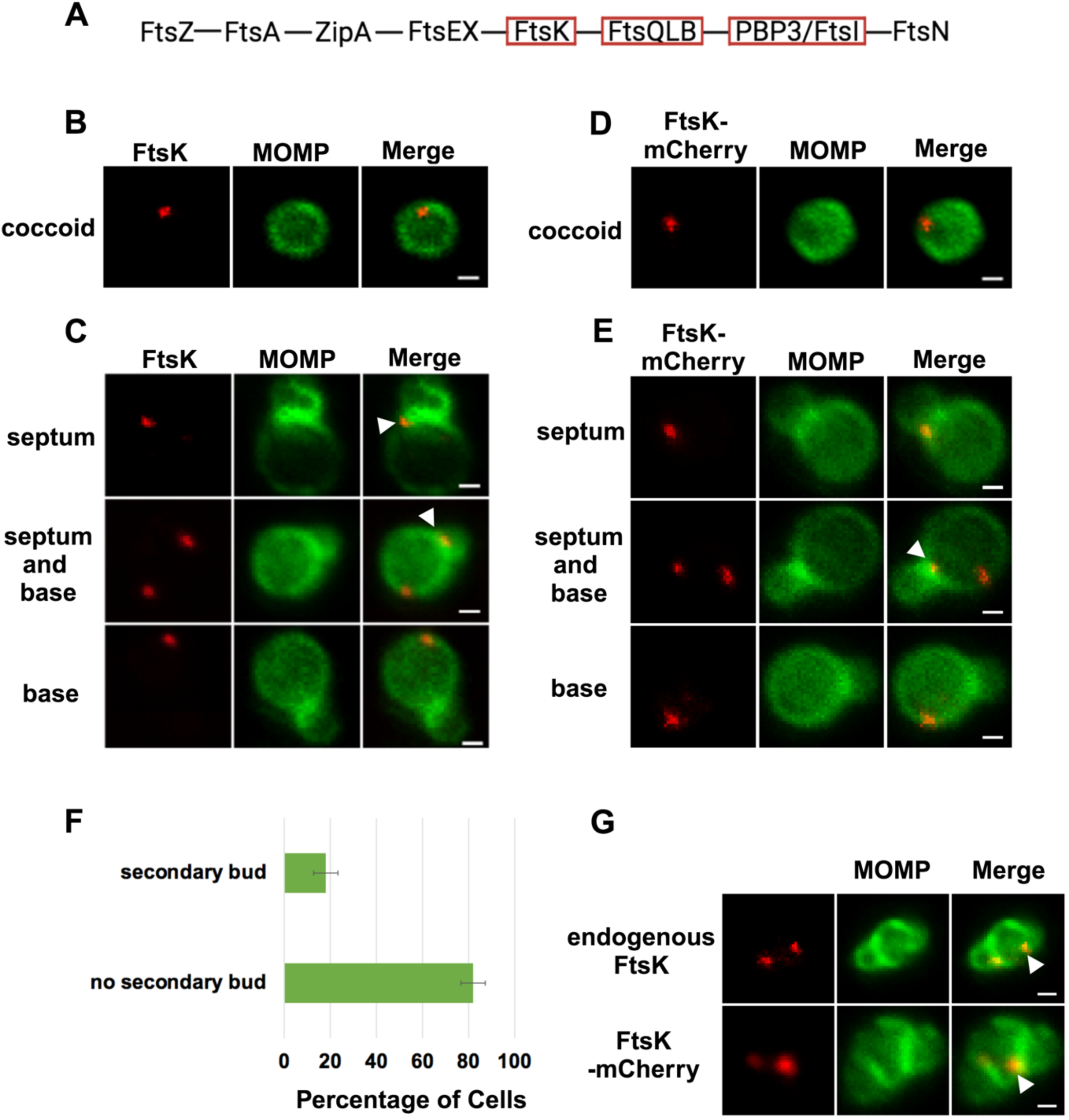
(A) The linear divisome assembly pathway of *E. coli* is shown. *Ct* encodes the divisome proteins boxed in red. HeLa cells were infected with *Ct* L2 and RBs were prepared at 21hpi. The cells were fixed and stained with MOMP (green) and FtsK (red) antibodies. The distribution of FtsK in (B) coccoid cells and in (C) cell division intermediates that had not initiated secondary bud formation is shown. Bars are 1 μm. HeLa cells were infected with *Ct* transformed with the pBOMB4 -Tet (-GFP) plasmid encoding FtsK-mCherry. The fusion was induced with 10nM aTc for 1 hr. and RBs were prepared from infected HeLa cells at 21hpi and stained with MOMP antibodies (green). The distribution of MOMP relative to the mCherry fluorescence (D) in coccoid cells and in (E) cell division intermediates that had not initiated secondary bud formation is shown. Bars are 1μm. Arrowheads in C and E denote foci of FtsK above or below the MOMP-stained septum. (F) HeLa cells were infected with *Ct* L2. At 21hpi, the cells were harvested and RBs were stained with MOMP antibodies. The number of dividing cells that had initiated secondary bud formation was quantified in 150 cells. Three independent replicates were performed, and the values shown are the average of the 3 replicates. (G) Endogenous FtsK and FtsK-mCherry accumulate in foci at the septum of secondary buds (marked with arrowheads).

To investigate the distribution of other putative chlamydial divisome components during budding, we transformed *Ct* with plasmids encoding PBP2, PBP3, or MreC with an N-terminal mCherry tag. IFU assays demonstrated that the aTc induced overexpression of the PBP2, PBP3, and MreC fusions had no effect on the developmental cycle progression of *C*t (Supp. Fig. S2A). In addition, blotting analyses revealed that mCherry antibodies primarily detected single species with the predicted molecular mass of the FtsK, PBP2, PBP3, and MreC fusions in lysates prepared from induced cells (Supp. Fig. S1B). The PBP2, PBP3, and MreC fusions were induced by the addition of 10nM aTc to infected cells at 19hpi, and the induced cells were harvested at 21hpi and stained with MOMP antibodies. Imaging analyses revealed that the PBP2, PBP3, and MreC fusions accumulated in foci in coccoid cells (Fig. 2A), and in cell division intermediates, the fusions accumulated in foci at the septum, in foci at the septum and at the base of the progenitor mother cell, or in foci only at the base of the progenitor mother cell (Fig. 2B). Similar analyses with an MreB_6xHis fusion (Lee 2020) revealed that MreB exhibited a similar localization profile (Figs. 2A and B). Each of these fusions, like FtsK, were restricted to one side and were often slightly above or slightly below (marked with arrowheads in Fig. 2B) the MOMP-stained septum in dividing cells. The foci of the fusions were also detected at the base of secondary buds (arrowheads in Supp. Fig. S2B). Quantification of the localization profiles of endogenous FtsK and the various fusion proteins revealed that the distribution profile of FtsK-mCherry accurately reflected the distribution of endogenous FtsK (Fig. 2C). Furthermore, a greater percentage of FtsK was associated with the base of dividing cells (including cells with septum and base, and cells with base alone) suggesting that FtsK associates with nascent divisomes at the base of dividing cells prior to the other putative divisome proteins assayed. Finally, this analysis suggested that MreB associated with nascent divisomes at the base of dividing cells after mCherry-PBP2 and mCherry-PBP3 (marked with # in Fig. 2C). The localization profiles of the chlamydial divisome proteins (Figs. 1 and 2) likely reflect the assembly of divisome complexes at the septum and at the base of the progenitor mother cell, and the disassembly of the septal divisome when divisome proteins are only present at the base of the mother cell.

**Figure 2:**
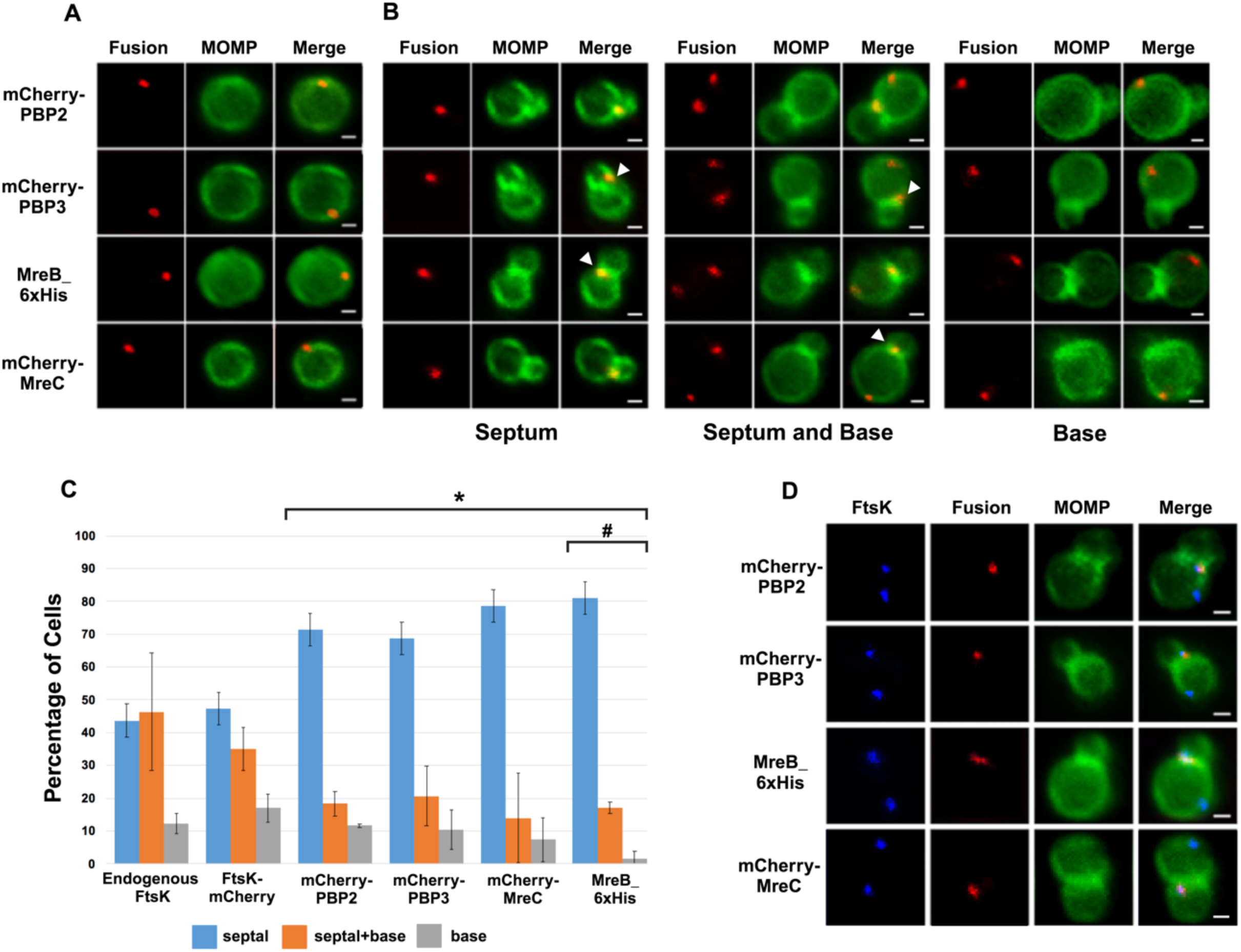
HeLa cells were infected with *Ct* transformed with PBP2, PBP3, or MreC with an N-terminal mCherry tag, or with *Ct* transformed with an MreB_6xHis fusion (Lee 2020). Each of the fusions was induced by adding 10nM aTc to the media at 17hpi. Lysates were prepared at 21hpi and the cells were fixed and stained with a MOMP antibody. The distribution of the mCherry fluorescence in (A) coccoid cells and in (B) dividing cells that had not initiated secondary bud formation is shown. Cells expressing the MreB_6xHis fusion were stained with rabbit anti-6x His antibody (red) and MOMP antibodies (green). Dividing cells with foci at the septum, foci at the septum and foci at the base of the mother cell, or foci at the base alone are shown for each of the fusions. Arrowheads in B denote foci of the divisome proteins above or below the plane of the MOMP-stained septum. (C) HeLa cells were infected with *Ct* L2. Alternatively, cells were infected with *Ct* that were transformed with FtsK-mCherry, mCherry-PBP2, mCherry-PBP3, mCherry-MreC, or MreB_6xHis. The cells expressing the fusions were induced with aTc for 1.5hrs then fixed at 21hpi and the distribution of endogenous FtsK, or the mCherry fluorescence in cells inducibly expressing the mCherry fusions, or the distribution of MreB in cells where the MreB_6xHis fusion was inducibly expressed was quantified. A small fraction (2 %) of dividing cells contained multiple foci of each of the divisome proteins at the septum and/or the base of the cell. Since these cells were relatively rare, we included these cells with cells that contained a single foci in the quantification. The localization profiles were quantified in 100 cells. Three independent replicates were performed, and the values shown are the average of the 3 replicates. Chi-squared analysis revealed that the localization profiles of endogenous FtsK and FtsK-mCherry are not statistically different from each other, but they are statistically different than the PBP2, PBP3, MreC and MreB localization profiles (* – p<0.009). The localization profile of the MreB fusion is also statistically different than the localization profiles of the mCherry fusions of PBP2 and PBP3 (#-p = 0.05). (D) Hela cells were infected with *Ct* transformed with PBP2, PBP3, or MreC with a N-terminal mCherry tag, or with *Ct* transformed with an MreB_6xHis fusion (Lee 2020). The fusions were induced by adding 10 nM aTc to the media at 17hpi. The cells were harvested at 21hpi and *Ct* were harvested and stained with FtsK and MOMP antibodies. The cells expressing the MreB fusion were stained with FtsK, MOMP, and 6xHis antibodies. Imaging analyses revealed that FtsK was present in foci at the septum and in foci at the base in these cells, while each of the fusions was only detected at the septum where they overlapped the distribution of septal FtsK (Bars are1μm).

Since it was possible that the localization profiles of the mCherry-PBP2 and mCherry-PBP3 fusions were at least in part due to their induced over-expression, we performed similar studies with rabbit antibodies generated against peptides derived from PBP2 or PBP3 (Ouellette SP 2012). Blotting analyses (Supp. Fig. S1C) with these antibodies revealed that they recognized mCherry-PBP2 and mCherry-PBP3 in *Ct* lysates, and immunofluorescent staining with the PBP2 and PBP3-specific antibodies (Supp. Fig. S1D) completely overlapped the mCherry fluorescence in cells when the mCherry PBP2 and PBP3 fusions were inducibly expressed in *Ct*. Imaging analyses with the antibodies that recognize endogenous PBP2 and endogenous PBP3 indicated that these antisera detected foci in coccoid cells, and in cell division intermediates, the PBP2 and PBP3 antibodies detected foci at the septum, foci at the septum and at the base of the mother cell, or foci at the base alone (Supp. Fig. S3A). Quantification revealed that the localization profiles of endogenous PBP2 and PBP3 in division intermediates (Supp. Fig. S3B) were not statistically different than the localization profiles observed for the mCherry fusions of PBP2 and PBP3 (Fig. 2C).

The quantification in Fig. 2C suggested that FtsK is recruited to nascent divisomes that form at the base of dividing cells prior to the other divisome components assayed. This hypothesis was tested by staining cells expressing the PBP2, PBP3, MreC, or MreB fusions with antibodies that recognize endogenous FtsK. Imaging analyses revealed that in a subset of cells, FtsK was detected in foci at the septum and at the base of dividing cells, while each of the fusions was only detected at the septum where they overlapped the distribution of septal FtsK (Fig. 2D), indicating that FtsK is recruited to nascent divisomes at the base of the cell prior to the other divisome components assayed. Additional analyses revealed that FtsK also overlapped the distribution of the PBP2, PBP3, MreC, and MreB fusions following their appearance at the base of dividing cells (Supp. Fig. S4A). Quantification revealed the percentage of cells in which FtsK overlapped the distribution of each of the fusions at the septum, at the septum and the base, and at the base alone in division intermediates (Supp. Fig. S4B).

### MreB Filament Formation is not Required for Foci Formation by FtsK, PBP2, and PBP3

MreB was one of the last components that associated with nascent divisomes forming at the base of the progenitor mother cell (Fig. 2C). To investigate whether MreB filament formation was required for the formation of foci by the other chlamydial divisome components, HeLa cells were infected with *Ct* transformed with the FtsK, PBP2, PBP3, MreC, or MreB fusions, and the fusions were induced by adding 10nM aTc to the media of the infected cells at 20hpi for 1hr. During the induction period, cells were incubated in the absence (Figs. 3B and 3D) or presence (Figs. 3C and 3E) of the MreB inhibitor, A22, which inhibits MreB filament formation (Bean, 2009). RBs were harvested at 21hpi and stained with the appropriate antibodies to assess the effect of A22 on the cellular distribution of the fusions. As previously shown (Ouellette SP 2012; Cox 2020), A22 inhibits chlamydial budding and most cells in the population were coccoid following A22 treatment (Fig. 3A). Furthermore, approximately 50% of the untreated control cells were coccoid, which is consistent with prior estimates of the number of non-dividing RBs at this stage of the developmental cycle (Lee 2018), indicating that our lysis procedure does not lead to a bias in the number of non-dividing coccoid cells in the population. MreB in coccoid cells adopted a diffuse pattern of localization following A22 treatment (Fig 3). A22 also had a statistically significant effect on the percent of coccoid cells containing MreC foci, but it did not affect the ability of FtsK, PBP2, or PBP3 to form foci in coccoid cells (Figs. 3D and 3E). These data indicate that MreB filaments do not function as a scaffold that is necessary for the assembly of all divisome components in *Ct*.

**Figure 3:**
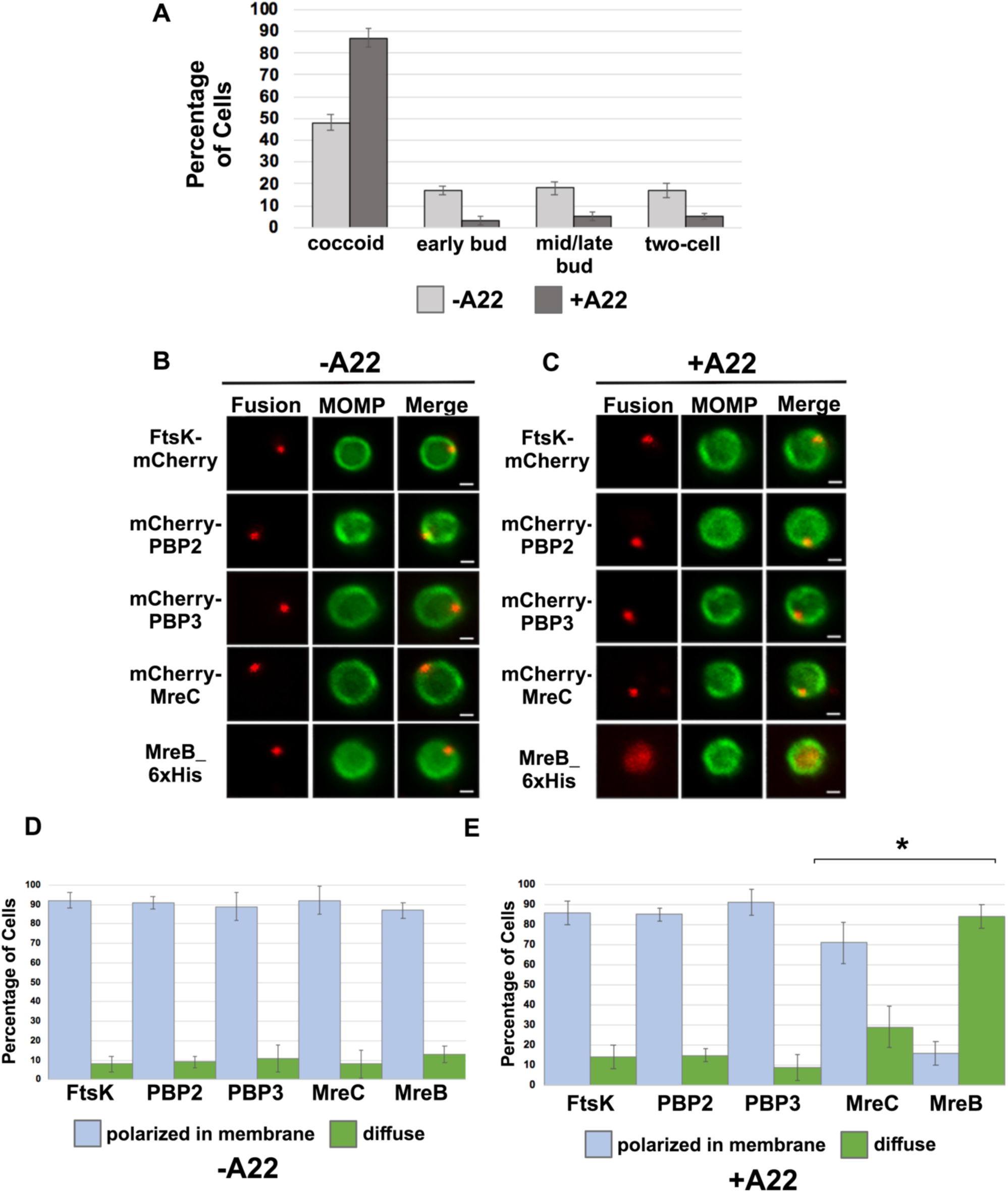
(A) HeLa cells infected with *Ct* L2 were treated with 75*μ*M A22 for 1 hour. Control cells were not treated with A22. Lysates were prepared at 21hpi and the number of coccoid and dividing cells in the population were quantified in 100 cells. Three independent replicates were performed, and the values shown are the average of the 3 replicates. (B-E) Alternatively, HeLa cells were infected with *Ct* transformed with plasmids encoding FtsK-mCherry, mCherry-PBP2, mCherry-PBP3, mCherry-MreC, or MreB-6xHis. The fusions were induced at 20hpi with 10nM aTc for 1hr in the absence (B and D) or presence (C and E) of 75μM A22. Coccoid cells prepared from the infected cells at 21hpi were stained with MOMP antibodies (green). The MreB-6xHis fusion was also stained with 6xHis antibodies (red). Panel B shows the distribution of the fusions in untreated coccoid cells. Panel C illustrates the effect of A22 on the localization of the fusions in coccoid cells. Bars in B and C are 1μm. The distribution of FtsK-mCherry, mCherry-PBP2, mCherry-PBP3, mCherry-MreC, and MreB-6xHis was quantified in (D) control coccoid cells and in (E) A22-treated cells coccoid cells (n=50) is shown. Three replicates were performed, and the values shown in D and E are the averages of the 3 replicates. Student T-test indicated that A22 had a statistically significant effect on the localization of MreB and MreC (* – p<0.01).

### Effect of *ftsk* and *pbp2* Knockdown on Cell Division and Divisome Assembly in *Ct*

To further investigate the mechanisms that regulate divisome assembly in *Ct*, we inducibly repressed the expression of *ftsK* or *pbp2* using CRISPRi technology, which has been used to inducibly repress the expression of genes in *Ct* (Ouellette SP 2021). CRISPRi employs a constitutively expressed crRNA that targets an inducible dCas enzyme (dCas12) to specific genes where it binds but fails to cut, thus inhibiting transcription. We transformed *Ct* with the pBOMBL12CRia plasmid that constitutively expresses an *ftsK* or *pbp2*-specific crRNA, which target sequences in the *ftsK* or *pbp2* promoter regions. To determine whether *ftsK* and *pbp2* transcript levels were altered using this CRISPRi approach, dCas12 expression was induced by the addition of 5nM aTc to the media of infected cells at 8hpi. Control cells were not induced. Nucleic acids were isolated from induced cells and uninduced control cells at various times, and RT-qPCR was used to measure *ftsK* or *pbp2* transcript levels. This analysis revealed that the induction of dCas12 resulted in ∼10-fold reduction in *ftsK* transcript levels by 15hpi in cells expressing the *ftsK-*targeting crRNA (Supp. Fig. S5A), and ∼8-fold reduction in *pbp2* transcript levels in cells expressing the *pbp2*-targeting crRNA (Supp. Fig. S5B), while these crRNAs had minimal or no effect on chlamydial *euo* and *omcB* transcript levels, suggesting that the *ftsK* and *pbp2* crRNAs specifically inhibit the transcription of *ftsK* and *pbp2* (Supp. Figs. S5A and S5B). To investigate the effect of *ftsK* or *pbp2* down-regulation on developmental cycle progression, dCas12 was induced by the addition of aTc to the media of infected cells at 4hpi. Control cells were not induced. The cells were then fixed at 24hpi and stained with MOMP and Cas12 antibodies. Imaging analysis revealed that *Ct* morphology was normal and dCas12 was undetectable in the inclusions of uninduced control cells, while foci of dCas12 were observed in the induced cells, and *Ct* in the inclusion exhibited an enlarged aberrant morphology (Supp. Figs. S5C and S5D), suggesting that the inducible knockdown of *ftsK* or *pbp2* blocks chlamydial cell division. In additional studies, we induced dCas12 at 17hpi in cells expressing the *ftsK* or *pbp2*-targeting crRNAs. Lysates were prepared and the cells were fixed at 21hpi, and localization studies revealed that foci of endogenous FtsK and PBP2 were almost undetectable when *ftsK* or *pbp2* were transiently knocked down using this CRISPRi approach (Supp. Fig. S5E).

To assess whether the knockdown of *ftsK* or *pbp2* arrests *Ct* division at a specific stage of polarized budding, HeLa cells were infected with *Ct* transformed with the pBOMBL12CRia plasmids encoding the *ftsK* or *pbp2*-targeting crRNAs. At 17hpi, dCas12 was induced by the addition of 20nM aTc to the media. Control cells were not induced. RBs were harvested from induced and uninduced control cells at 22hpi and stained with MOMP antibodies, and imaging analyses quantified the division intermediates present in the population. These analyses revealed that >60% of the *Ct* in the uninduced controls were at various stages of polarized budding (Figs. 4A and B), while ∼90% of the cells were coccoid when *ftsK* was knocked down (Fig. 4A), and ∼85% of the cells were coccoid following *pbp2* knockdown (Fig. 4B), suggesting that the initiation of polarized budding of *Ct* is inhibited when *ftsK* or *pbp2* are knocked down.

**Figure 4:**
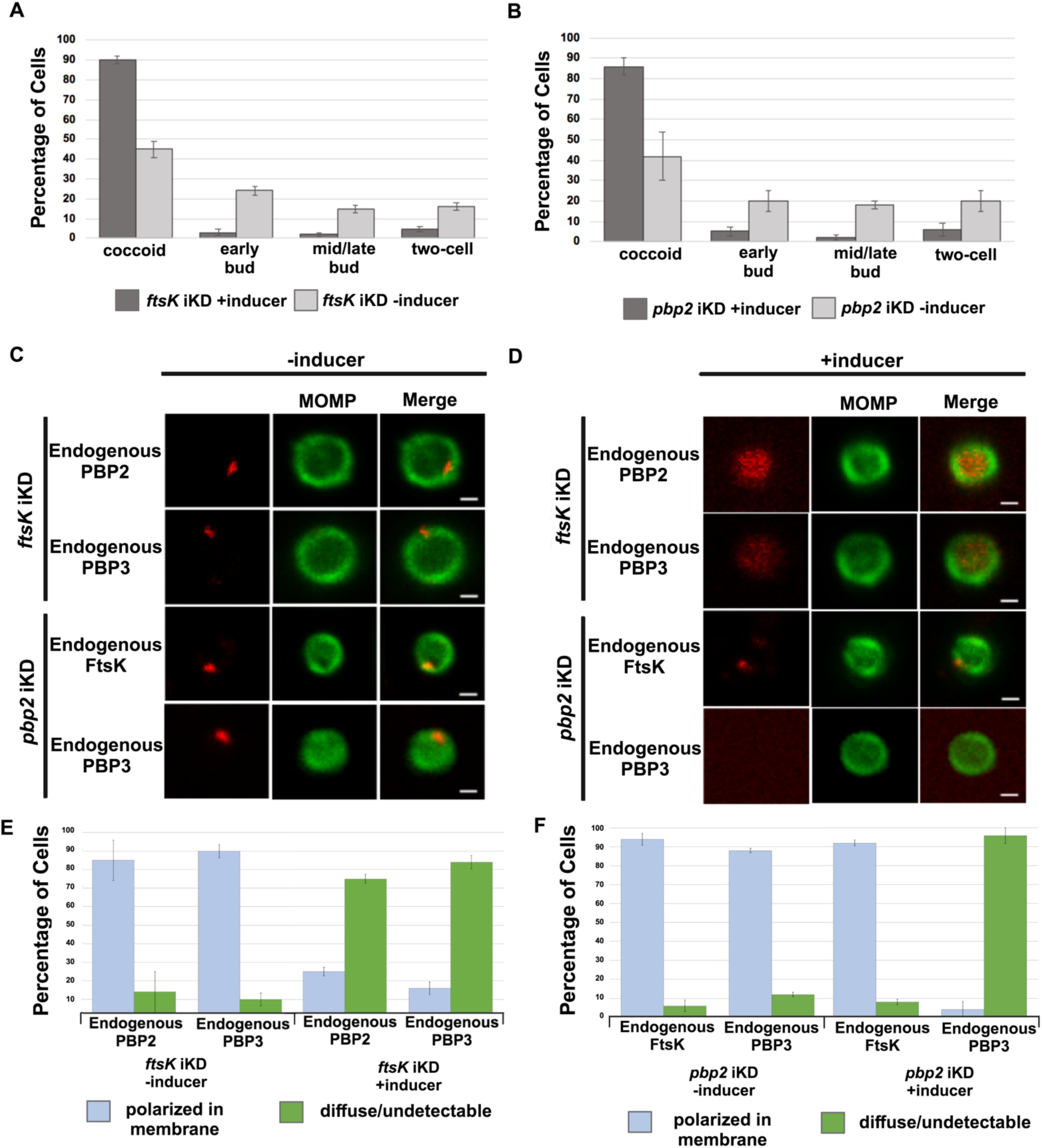
HeLa cells were infected with *Ct* transformed with the pBOMBL-12CRia plasmid, which constitutively expresses a *ftsK* crRNA or *pbp2* crRNA and encodes dCas12 under the control of an aTc-inducible promoter. dCas12 was induced at 17hpi by adding 5nM aTc to the media. In a control infection, the expression of dCas12 was not induced. Cells were harvested at 24hpi and the morphology of *Ct* in induced and uninduced control cells was assessed in 250 cells. 3 replicates were performed, and the values shown are the averages of the 3 replicates (A and B). The localization of endogenous FtsK, endogenous PBP2, and endogenous PBP3 was assessed in cells transformed with the pBOMBL-12CRia plasmid that targets *ftsK* or *pbp2.* The localization is shown in coccoid cells where dCas12 expression was (C) uninduced or (D) induced. White bars are 1μm. (E and F) The localization profiles of FtsK, PBP2, and PBP3 were quantified in uninduced and induced cells. 3 replicates were performed, and the values shown are the averages of the 3 replicates.

We next examined whether the knockdown of *ftsK* affected foci formation by PBP2 or PBP3 in coccoid cells. HeLa cells were infected with *Ct* transformed with the pBOMBL12CRia plasmid encoding the *ftsK*-targeting crRNA. At 17hpi, dCas12 was induced by the addition of 10nM aTc to the media. Control cells were not induced. RBs were harvested from induced and uninduced control cells at 22hpi, and the localization of endogenous PBP2 and PBP3 in coccoid cells was assessed (Figs. 4C and 4D). This analysis revealed that the number of coccoid cells with polarized foci of PBP2 and PBP3 were reduced by approximately 80% following *ftsK* knockdown (Fig. 4E). In similar analyses, we assessed the effect of *pbp2* knockdown on the ability of FtsK and PBP3 to form foci in coccoid cells. While FtsK retained its ability to form foci in coccoid cells following *pbp2* knockdown, foci of PBP3 were almost entirely absent in *pbp2* knockdown cells (Figs. 4C and 4D). Quantification of these assays revealed that FtsK is necessary for foci formation by both the PBP2 and PBP3 transpeptidases, while PBP2 is necessary for foci formation by PBP3 (Figs. 4E and 4F). Our data place FtsK upstream of, and necessary for, the addition of PBP2 and PBP3 to the *Ct* divisome, and PBP2 upstream of and necessary for the addition of PBP3 to the *Ct* divisome. These results are consistent with inhibitor studies that indicated PBP2 acts upstream of PBP3 in the polarized budding process of *Ct* (Cox 2020).

To investigate whether the catalytic activity of PBP2 is necessary to maintain its association with the *Ct* divisome, HeLa cells were infected with *Ct*, and mecillinam, an inhibitor of the transpeptidase activity of PBP2 (Kocaoglu 2015; Cox 2020), was added to the media of infected cells at 20hpi. Cells incubated in the absence of mecillinam were included as a control. Mecillinam-treated and control cells were harvested at 22hpi and the effect of inhibiting the catalytic activity of PBP2 on the localization of FtsK, PBP2, and PBP3 was determined. As shown previously, mecillinam blocks chlamydial division (Ouellette SP 2012; Cox 2020), and most cells in the population assumed a coccoid morphology (Fig. 5A). We then determined the localization of endogenous FtsK, PBP2, and PBP3 in drug-treated and control coccoid cells. Mecillinam treatment resulted in a ∼50% reduction in the number of cells with polarized foci of PBP2 (Figs. 5B - 5E). There was a similar reduction in polarized foci of PBP3 following mecillinam treatment (Figs. 5B - 5E). These data indicate that the catalytic activity of PBP2 is necessary for PBP2 to efficiently associate with or maintain its association with polarized divisome complexes. Furthermore, consistent with *pbp2* knockdown studies, PBP3 association with the divisome complex is dependent on the prior addition of PBP2 to the complex, but foci formation by FtsK is unaffected when PBP2 foci are reduced in number (Fig. 5).

**Figure 5:**
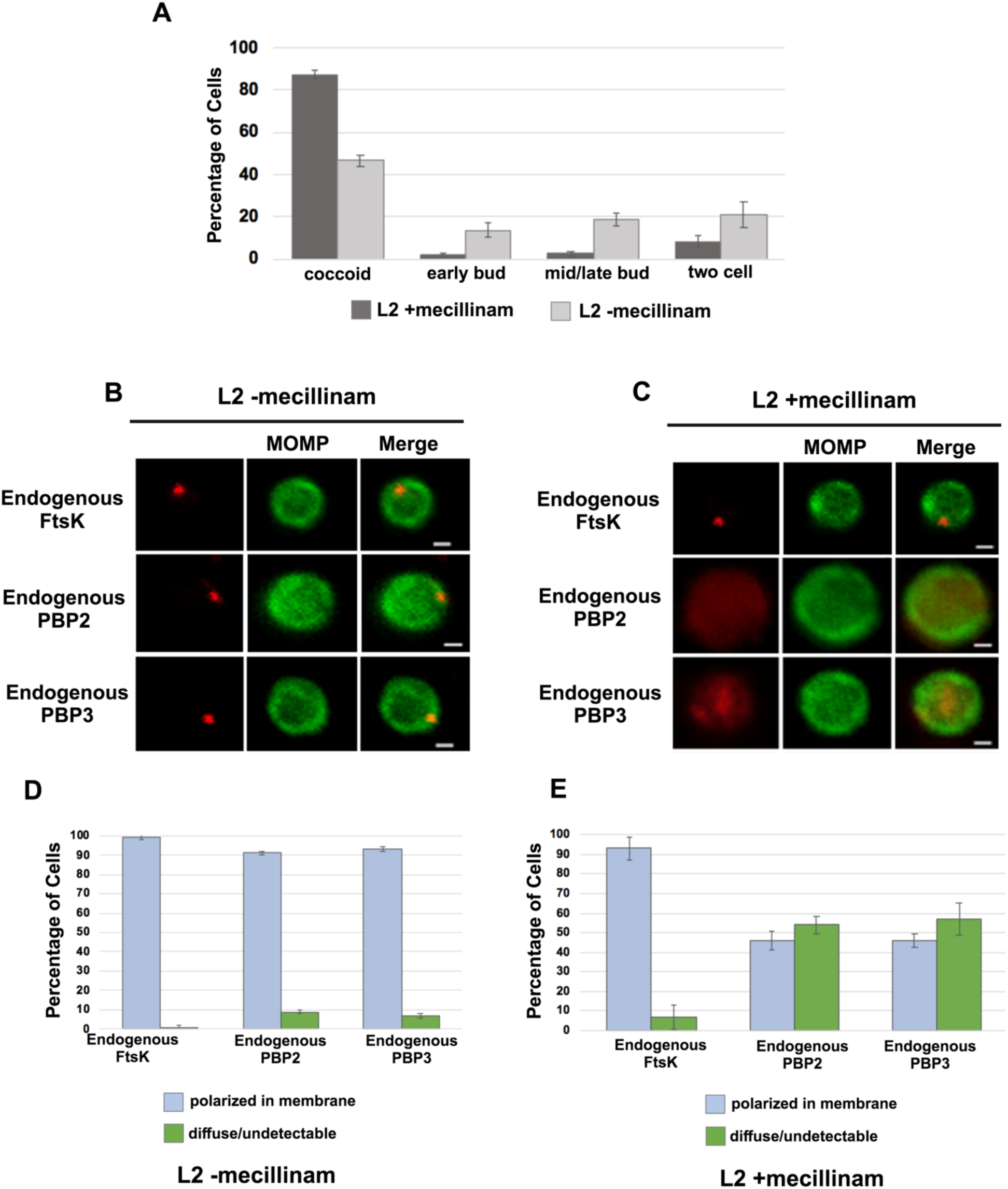
Effect of mecillinam on endogenous FtsK, PBP2, and PBP3 localization. (A) HeLa cells were infected with *Ct* and 20μM mecillinam was added to the media at 17hpi. Untreated coccoid cells were included as a control. The cells were harvested at 21hpi and the morphology of MOMP-stained cell was assessed in 200 cells. 3 replicates were performed, and the values shown are the averages of the 3 replicates. (B and C) The localization of endogenous FtsK, endogenous PBP2, and endogenous PBP3 in untreated coccoid or in mecillinam-treated coccoid cells is shown. Bars are1μM. (D and E) Localization of FtsK, PBP2, and PBP3 in untreated and mecillinam-treated coccoid cells was quantified in 50 cells. Three replicates were performed, and the values shown are the averages of the 3 replicates.

### Effect of Inhibitors and *ftsk* Knockdown on PG Organization in *Ct*

To assess the morphology of PG at the septum and the base of dividing cells, we used an EDA-DA labeling strategy (Liechti 2014; Cox 2020). This approach enabled the detection of PG foci, bars, and rings in dividing *Ct* (Liechti 2021). Our imaging analysis revealed that in some instances PG foci were detected at both the septum and at the base of the progenitor mother cell (Fig. 6A), and in other instances PG organization at the septum and the base differed. In the example shown (Fig. 6B), a PG foci was detected at the septum while a PG ring was detected at the base of the mother cell. In additional analyses, we compared the localization of mCherry-PBP3 to the localization of PG in cells where the expression of this mCherry fusion had been induced by the addition of aTc to the media. This analysis, which was restricted to PG formation at the septum of dividing cells, revealed that multiple foci of PBP3 were associated with a septal PG ring (Fig. 6C). Furthermore, the PG ring was at an angle relative to the MOMP-stained septum. Similar analyses revealed that 2 foci of endogenous FtsK were associated with PG that was again at an angle relative to the MOMP-stained septum. This was true even though the PG had not fully reorganized into a ring structure (Fig. 6C).

**Figure 6:**
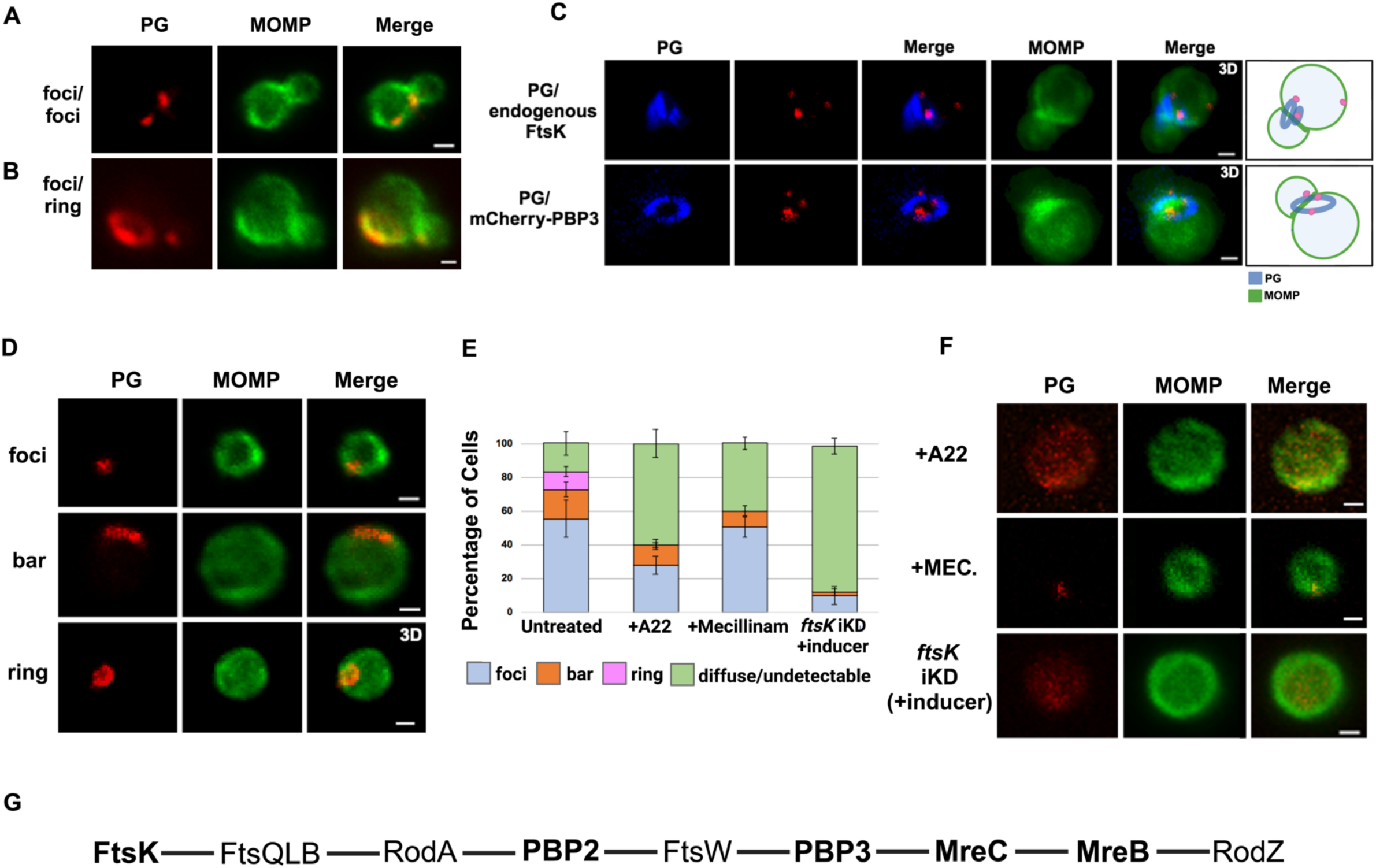
PG distribution in *Ct*. HeLa cells were infected with *Ct* L2. At 17hpi, 4mM ethylene-DA-DA (EDA-DA) was added to the media, the cells were harvested at 21hpi, and the EDA-DA was click labeled and compared to the distribution of MOMP. (A) Imaging analyses revealed that PG formed foci at the septum and at the base in some dividing cells. (B) Imaging analyses revealed that PG organization at the septum and at the base of some dividing cells differed. In this example, a PG foci was detected at the septum and a PG ring was detected at the base of a dividing cell. (C) The localization of click-labeled PG was compared to the localization of endogenous FtsK and mCherry-PBP3 at the septum of dividing cells. 3D projections revealed that multiple foci of each protein are associated with PG intermediates. Cartoons are included to assist the reader in visualizing the angled orientation of PG relative to the MOMP-stained septum in the dividing cells. (D) PG organization in untreated coccoid cells. (E) Quantification of PG organization in untreated coccoid cells, A22-treated coccoid cells, mecillinam-treated coccoid cells, and in coccoid cells resulting from the inducible knockdown of *ftsk*. Fifty cells were counted for each condition. Three replicates were performed and the average from the 3 replicates is shown. (F) PG organization in A22-treated and mecillinam-treated coccoid cells, and in coccoid cells resulting from the inducible knockdown of *ftsk* is shown. Bars are 1μm. (G) Putative *Ct* divisome assembly pathway is shown. Proteins characterized in this study are bolded. The ordering of the remaining proteins is based on the assembly of the divisome and elongasome in *E. col*i (Du 2017; Liu 2020).

We then determined the effect of A22 and mecillinam on PG synthesis/organization. Since both of these drugs induce *Ct* to assume a coccoid morphology, we initially characterized PG organization in untreated coccoid cells. We detected foci, bars, or rings in ∼80% of untreated coccoid cells (Figs. 6D and 6E), which make up ∼50% of the cells in the inclusion at this stage of the developmental cycle (Lee 2018). Furthermore, each of these PG intermediates exhibited a polarized distribution in untreated coccoid cells (Fig. 6D). Although we cannot rule out that continued PG synthesis and reorganization occurs in polarized division intermediates, PG rings can arise prior to any of the morphological changes that occur during the polarized division of *Ct*.

Prior studies have shown that inhibitors of MreB filament formation prevent the appearance of PG-containing structures in *Ct* (Liechti 2014; Ouellette SP 2022). To assess whether MreB filament formation is required for PG synthesis and/or PG reorganization, we infected HeLa cells with *Ct*, and EDA-DA and A22 were added to the media of infected cells at 18hpi. The cells were harvested at 22hpi, lysates were prepared, and PG localization was determined. These analyses revealed that PG was diffuse/undetectable in the majority of A22-treated cells, and, in those cells where PG was still detected, it could not convert into ring structures (Figs. 6E and 6F). In similar experiments, we assessed the effect of mecillinam on the appearance of PG intermediates in coccoid cells. These analyses revealed that PG formed discrete foci or bars in 60% of mecillinam-treated cells (Figs. 6E and 6F). However, these PG intermediates could not convert into PG rings when the transpeptidase activity of PBP2 was inhibited. Finally, we assessed PG organization in cells where *ftsK* was knocked down by inducing dCas12 in the *ftsK* knockdown strain by the addition of aTc to the media of infected cells at 17hpi. Cells were fixed at 21hpi, and localization studies revealed that *ftsK* knockdown had the most dramatic effect on PG localization, which was diffuse/undetectable in ∼90% of the cells assayed. Inhibiting divisome assembly by knocking down *ftsK* almost entirely prevented the accumulation of all PG-containing intermediates in *Ct* (Figs. 6E and 6F).

## Discussion

The results presented here provide insight into the molecular mechanisms governing the FtsZ-less polarized cell division process of *Ct*. This study is the first to document the ordered assembly of divisome proteins in *Ct* and to investigate the roles of divisome proteins in regulating PG synthesis/organization in this obligate intracellular bacterial pathogen. Our studies showed that the divisome in *Ct* is hybrid in composition, containing elements of the divisome and elongasome from other bacteria, and a putative pathway for the assembly of the novel hybrid divisome of *Ct* is shown in Fig. 6G. Although we focused our study on a subset of the divisome and elongasome proteins that *Chlamydia* expresses (bolded in Fig. 6G), our results support our conclusion that chlamydial budding is dependent upon a hybrid divisome complex and that FtsK is required for the assembly of this hybrid divisome. At this time, we cannot rule out that other proteins act upstream of FtsK to initiate divisome assembly in this obligate intracellular bacterial pathogen.

The domain organization of FtsK is highly conserved among bacteria. The C-terminus of the protein mediates the ATP-dependent directional translocation of the protein along DNA through interaction with KOPS (Fts**K o**rienting **p**olar **s**equences) (Barre, 2001; Massey, 2006) and stimulates XerCD-dependent recombination at the *dif* site near the chromosomal terminus to decatenate chromosomal dimers that arise from homologous recombination during DNA replication to monomers (Blakely, 1993; Massey, 2006). The C-terminus of chlamydial FtsK is 41% identical to the C-terminus of *E. coli* FtsK. The N-terminus of *E. coli* FtsK encodes 4 predicted transmembrane spanning helices, which are sufficient to direct the protein to the septum in dividing cells (Yu, 1998). The N-terminus of chlamydial FtsK, which is also predicted to encode 4 transmembrane-spanning helices, is 34% identical to *E. coli* FtsK.

Our analyses revealed a novel spatiotemporal localization pattern of FtsK during the chlamydial division process. Chlamydial FtsK forms discrete foci at the septum, foci at the septum and at the base of the mother cell, or in foci only at the base of the mother cell (Figure 1). Our data indicate that the foci at the base of the mother cell correspond to nascent divisome complexes that form prior to the formation of a secondary bud at the base of the progenitor mother cell. Our analyses further revealed there was no correlation between the stage of bud formation by the initial bud (early, mid-late; Ouellette SP, 2022), and the appearance of nascent divisomes at the base of the progenitor mother cell (data not shown).

The *Ct* divisome is hybrid in nature, containing elements of the divisome (FtsK and PBP3) and elongasome (PBP2, MreB, and MreC) from other bacteria. Each of these proteins formed foci at the septum, foci at the septum and at the base of the mother cell, or foci only at the base of the mother cell, and the foci of each protein were restricted to one side of the MOMP-stained septum (Figs. 1 and 2). Knockdown of *ftsK* using CRISPRi revealed that FtsK is necessary for the assembly of this hybrid divisome complex (Fig. 4). Knockdown and inhibitor studies further revealed that PBP2 is necessary for the addition of PBP3 to the divisome in *Ct* (Figs. 4 and 5).

The chlamydial divisome proteins all form foci in coccoid cells (Figs. 1 and 2), and FtsK forms foci in dividing *Ct* that only partially overlap the distribution of PBP2, PBP3, MreC, and MreB (Fig. 2 and Supp. Fig. S4A). The lack of overlap of FtsK with MreB and MreC is most clearly evident in Supp. Fig. S4A, and while it is unclear why FtsK only partially overlaps the distribution of the other divisome components in dividing *Ct*, MreB (Liechti 2014; Kemege 2015; Lee 2020) and MreC (Supp. Fig. S6A) can reorganize into rings in dividing cells, and the MreC rings we detected, like PG rings (Fig. 6), were at an angle relative to the MOMP-stained septum. MreC also forms rings in coccoid cells (Supp. Fig. S6B). Although MreB and MreC form rings in *Ct*, we never detected FtsK rings in dividing or coccoid chlamydial cells. The relevance of the angled orientation of PG and MreC rings relative to the MOMP-stained septum in division intermediates is unclear. However, it appears to be a conserved feature of the cell division process and may arise because the divisome proteins are often positioned slightly above or below the plane of the MOMP-stained septum (Figs. 1 and 2).

Previous studies hypothesized that MreB filaments may substitute for FtsZ and form a scaffold necessary for the assembly of the divisome in *Ct* (Ouellette SP 2012; Ouellette SP 2015; Ouellette SP 2020). However, our analyses have indicated that MreB is one of the last components recruited to nascent divisomes that form at the base of the mother cell in *Ct,* and localization studies revealed that foci formation by FtsK, PBP2, and PBP3 are not dependent on MreB filament formation (Fig. 3). Although our data indicate that MreB filaments do not form a scaffold necessary for the assembly of all components of the divisome in *Ct*, MreB filaments are necessary for the conversion of PG foci into PG rings in *Ct* (Fig. 6).

FtsZ treadmilling drives its rotational movement at the septum and this may be required for the positioning of peptidoglycan biosynthetic enzymes at the division plane in gram-negative and gram-positive bacteria (Bisson-Filho 2017; Yang 2017). The knockdown studies presented here demonstrated that in the absence of FtsZ, chlamydial FtsK is critical for divisome assembly and PG synthesis in *Ct,* and future real-time imaging studies will determine whether changes in the organization/distribution of FtsK occur during chromosomal translocation and/or cell division in *Chlamydia*. Furthermore, future studies will investigate the mechanisms that regulate the site of assembly of nascent divisomes in the mother cell during the polarized cell division process.

*Ct* is a member of the Planctomycetes/Verrucomicrobia/Chlamydia superphylum and members of the Chlamydia and Planctomycetes phyla do not encode FtsZ (Rivas-Marín 2016). *Planctospirus limnophila* is a member of the Planctomycetes that divides by polarized budding, and recent knockout studies (Rivas-Marin 2023) indicated that FtsK is the only protein of the chlamydial divisome we characterized here that is essential for the growth of this free-living organism. These results suggest that multiple mechanisms of FtsZ-independent polarized budding have evolved in members of this superphyla. It will be of interest in future studies to determine whether other members of the Planctomycetes that bud (Wiegand 2020) divide using a divisome apparatus similar to *Ct*.

## Materials and Methods

### Cell Culture

HeLa cells (ATCC, Manassas, VA) were cultured in Dulbecco’s Modified Eagle Medium (DMEM; Invitrogen, Waltham, MA) containing 10% fetal bovine serum (FBS, Hyclone, Logan, UT) at 37°C in a humidified chamber with 5% CO2. HeLa cells were infected with *Ct* serovar L2 434/Bu in the same media. Infections of HeLa cells with chlamydial transformants were performed in DMEM containing 10% FBS and 0.36 U/mL penicillin G (Sigma-Aldrich).

### Cloning

The plasmids and primers used for generating mCherry fusions of FtsK, PBP2, PBP3, and MreC are listed in Supp. Table S1. The chlamydial *ftsK*, *pbp2, pbp3,* and *mreC* genes were amplified by PCR with Phusion DNA polymerase (NEB, Ipswich, MA) using 10 ng *C*. *trachomatis* serovar L2 genomic DNA as a template. The PCR products were purified using a PCR purification kit (Qiagen) and inserted into the pBOMB4-Tet (-GFP) plasmid, which confers resistance to β-lactam antibiotics. The plasmid was cut at the NotI (FtsK-mCherry) or the KpnI (mCherry-PBP2, mCherry-PBP3, mCherry-MreC) site, and the chlamydial genes were inserted into the cut plasmid using the HiFi DNA Assembly kit (NEB) according to the manufacturer’s instructions. The products of the HiFi reaction were transformed into NEB-5αI^q^ competent cells (NEB) and transformants were selected by growth on plates containing ampicillin. DNA from individual colonies was isolated using a mini-prep DNA isolation kit (Qiagen), and plasmids were initially characterized by restriction digestion to verify the inserts were the correct size. Clones containing inserts of the correct size were DNA sequenced prior to use.

### DNA and RNA purification and RT-qPCR

Total nucleic acids were extracted from HeLa cells infected with *Ct* plated in 6-well dishes as described previously (Ouellette 2015, Ouellette, Blay et al. 2021). For RNA isolation, cells were rinsed one time with PBS, then lysed with 1mL Trizol (Invitrogen) per well. Total RNA was extracted from the aqueous layer after mixing with 200μL per sample of chloroform following the manufacturer’s instructions. Total RNA was precipitated with isopropanol and treated with DNase (Ambion) according to the manufacturer’s guidelines prior to cDNA synthesis using SuperScript III (Invitrogen). For DNA, infected cells were rinsed one time with PBS, trypsinized and pelleted before resuspending each pellet in 500μL of PBS. Each sample was split in half, and genomic DNA was isolated from each duplicate sample using the DNeasy extraction kit (Qiagen) according to the manufacturer’s guidelines. Quantitative PCR was used to measure *C*. *trachomatis* genomic DNA (gDNA) levels using an *euo* primer set. 150ng of each sample was used in 25μL reactions using standard amplification cycles on a QuantStudio3 thermal cycler (Applied Biosystems) followed by a melting curve analysis. *ftsK, pbp2, euo*, and *omcB* transcript levels were determined by RT-qPCR using SYBR Green as described previously (Ouellette SP 2021) (see Supp. Table S2 for primers used for measuring gDNA levels and RT-qPCR). Transcript levels were normalized to genomes and expressed as ng cDNA/gDNA.

### Transformation of *Ct*

*Ct* was transformed as described previously (Wang 2011). Briefly, HeLa cells were plated in a 10cm plate at a density of 5 × 10^6^ cells the day before beginning the transformation procedure. *Ct* lacking its endogenous plasmid (-pL2) was incubated with 10μg of plasmid DNA in Tris-CaCl_2_ buffer (10 mM Tris-Cl pH 7.5, 50 mM CaCl_2_) for 30 min at room temperature. HeLa cells were trypsinized, washed with 8mL of 1x DPBS (Gibco), and pelleted. The pellet was resuspended in 300μL of the Tris-CaCl_2_ buffer. *Ct* was mixed with the HeLa cells and incubated at room temperature for an additional 20 min. The mixture was added to 10mL of DMEM containing 10% FBS and 10 μg/mL gentamicin and transferred to a 10cm plate. At 48hpi, the HeLa cells were harvested and *Ct* in the population were used to infect a new HeLa cell monolayer in media containing 0.36 U/ml of penicillin G to select for transformants. The plate was incubated at 37°C for 48 hours. These harvest and re-infection steps were repeated every 48hrs until inclusions were observed.

### Immunofluorescence Microscopy

HeLa cells were seeded in 10cm plates at a density of 5 × 10^6^ cells per well the day before the infection. *Ct* L2 or chlamydial strains transformed with plasmids encoding FtsK-mCherry, mCherry-PBP2, mCherry-PBP3, or mCherry-MreC or with plasmids that direct the constitutive expression of the crRNAs targeting the *pbp2* or *ftsK* promoters were used to infect HeLa cells in DMEM. For experiments with the transformants, aTc was added to the media of infected cells at the indicated concentration and time. At 21hpi, cells were detached from the 10cm plate by scraping and pelleted by centrifugation for 30 seconds. The pellet was resuspended in 1 mL of 0.1x PBS (Gibco) and transferred to a 2mL tube containing 0.5mm glass beads (ThermoFisher Scientific). Cells were vortexed for 3 mins. then centrifuged at 800rpm for 2 mins. in a microfuge. 20μLs of the supernatant was mixed with 20μLs of 2x fixing solution (6.4% formaldehyde and 0.044% glutaraldehyde) and incubated on a glass slide for 10 min at room temperature. Cells were washed with 3 times with PBS, and the cells were permeabilized by incubation with PBS containing 0.1% Triton X-100 for 1 min. Cells were washed with PBS two times. For experiments with *Ct* L2, the cells were incubated with a goat primary antibody against the major outer-membrane protein (MOMP; Meridian, Memphis, TN), and the mouse primary antibody that recognizes endogenous FtsK raised against recombinant CT739 protein (https://doi.org/10.1099/mic.0.047746-0), or with rabbit antibodies raised against peptides derived from PBP 2 or PBP3 (Ouellette SP 2012). Briefly, chlamydial antigens or peptides emulsified with Freund’s incomplete adjuvant were used to immunize animals via intramuscular injections three times with an interval of 2 weeks. Antisera were collected from the immunized animals 2 to 4 weeks after the final immunization as the primary antibodies. After the primary antibody labeling, the cells were then rinsed with PBS and incubated with donkey anti-goat IgG (Alexa 488) and donkey anti-mouse IgG (Alexa 594) or donkey-anti-rabbit IgG (Alexa 594) secondary antibodies (Invitrogen). Experiments in which we visualized the distribution of the various mCherry fusions, the localization of the mCherry fluorescence was compared to the distribution of MOMP. In some experiments, we determined the distribution of the MreB_6x His fusion by staining cells expressing the fusion with a rabbit anti-6x His antibody (Abcam, Cambridge, MA) and the goat anti-MOMP antibody followed by the appropriate secondary antibodies. Cells were imaged using Zeiss AxioImager2 microscope equipped with a 100x oil immersion PlanApochromat objective and a CCD camera. During image acquisition, 0.3μm xy-slices were collected that extended above and below the cell. The images were collected such that the brightest spot in the image was saturated. The images were deconvolved using the nearest neighbor algorithm in the Zeiss Axiovision 4.7 software. Deconvolved images were viewed and assembled using Zeiss Zen-Blue software. For each experiment, three independent replicates were performed, and the values shown for localization are the average of the 3 experiments. In some instances, 3D projections of the acquired xy slices were generated using the Zeiss Zen-Blue software.

### Peptidoglycan (PG) labeling

PG was labeled by incubating cells with 4mM ethylene-D-alanine-D-alanine (E-DA-DA) as described (Cox 2020). The incorporated E-DA-DA was fluorescently labeled using the Click & Go^TM^ labeling kit (Vector Laboratories). The distribution of fluorescently labeled PG was compared to the distribution of MOMP and endogenous FtsK or the distribution of mCherry-PBP3. Three independent replicates were performed, and the values shown are the average of the 3 experiments.

### Inclusion forming unit assay

HeLa cells were infected with *Ct* (-pL2) transformed with the pBOMB4 Tet (-GFP) plasmid encoding the indicated aTc-inducible gene. At 8hpi, aTc was added to the culture media at the indicated concentration. Control cells were not induced. At 48hpi, the HeLa cells were dislodged from the culture dishes by scraping and collected by centrifugation. The pellet was resuspended in 1 mL of 0.1x PBS (Gibco) and transferred to a 2mL tube containing 0.5mm glass bead tubes (ThermoFisher Scientific). Cells were vortexed for 3 min. followed by centrifugation at 800rpm for 2 min. The supernatants were mixed with an equal volume of a 2x sucrose-phosphate (2SP) solution (ref) and frozen at −80°C. At the time of the secondary infection, the chlamydiae were thawed on ice and vortexed. Cell debris was pelleted by centrifugation for 5 min at 1k x g at 4°C. The EBs in the resulting supernatant were serially diluted and used to infect a monolayer of HeLa cells in a 24-well plate. The secondary infections were allowed to grow at 37°C for 24 hrs before they were fixed and labeled for immunofluorescence microscopy by incubating with a goat anti-MOMP antibody followed by a secondary donkey anti-goat antibody (Alexa Fluor 594). The cells were rinsed in PBS and inclusions were imaged using an EVOS imaging system (Invitrogen). The number of inclusions were counted in 5 fields of view and averaged. Three independent replicates were performed, and the values from the replicates were averaged to determine the number of inclusion forming units. Chi-squared analysis was used to compare IFUs in induced and uninduced samples.

### Effect of A22 and mecillinam on the profile of division intermediates and on PG and divisome protein localization in *Ct*

HeLa cells were infected with *Ct* transformed with the pBOMB4-Tet (-GFP) plasmid encoding FtsK-mCherry, mCherry-PBP2, mCherry-PBP3, mCherry-MreC, or MreB-6xHis. The fusions were induced at 20hpi with 10nM aTc for 1hr in the absence or presence of 75 μM A22. At 22hpi, cells were harvested and prepared for staining as described above. Three independent replicates were performed, and the values shown for localization are the average of the 3 experiments. HeLa cells were infected with *Ct* L2 and 20μM mecillinam (Sigma) was added to the media of infected cells at 17 hpi. Control cells were untreated. At 22 hpi, infected cells were harvested and RBs were prepared and stained with MOMP, FtsK, PBP2 or PBP3 antibodies as described above. Alternatively, cells were incubated with 4mM EDA-DA at 17hpi in the presence or absence of 20μM mecillinam. The cells were harvested at 22hpi, and RBs were prepared and PG was click-labeled, and its distribution was visualized in MOMP-stained cells as described above. Three independent replicates were performed, and the values shown for localization are the average of the 3 experiments.

### Immunoblotting

HeLa cells infected with *Ct* L2 were harvested by scraping the infected cells from the plate at 24hpi. Uninfected HeLa cells were included as a control. The HeLa cells were pelleted by centrifugation, resuspended in SDS sample buffer and electrophoresed on a 10% SDS polyacrylamide gel. The gel was electrophoretically transferred to nitrocellulose (Schleicher and Schuell), and the filter was incubated with mouse polyclonal antibodies raised against chlamydial FtsK. The filter was rinsed and incubated with 800 donkey anti-mouse IgG secondary antibodies (LICOR, Lincoln, NE) and imaged using a LICOR Odyssey imaging system.

HeLa cells were infected with *Ct* transformed with plasmids that inducibly express FtsK-mCherry, mCherry-PBP2, mCherry-PBP3, or mCherry-MreC. The fusions were induced by the addition of 10nM aTc to the media of infected cells at 17hpi. The cells were harvested and pelleted at 21hpi. The cell pellet was resuspended in 1 mL of 0.1x PBS (Gibco) and transferred to a 2mL tube containing 0.5mm glass beads (ThermoFisher Scientific). Cells were vortexed for 3 min. followed by centrifugation at 800rpm for 2 min. The supernatant was collected and centrifuged for 3 min at 13,000 rpm and the pellet containing *Ct* was resuspended in TBS containing 1% TX-100, 1X protease inhibitor cocktail (Sigma), and 5μM lactacystin. The suspension was sonicated 3 times on ice and centrifuged at 13,000 rpm for 3 mins. The supernatant was collected and mixed with SDS sample buffer. The samples were boiled and electrophoresed on a 10% SDS polyacrylamide gel, and the gel was electrophoretically transferred to nitrocellulose. The blots from these analyses were probed with a rabbit anti-mCherry primary antibody (Invitrogen) and a 800 donkey anti-rabbit IgG secondary antibodies (LICOR, Lincoln, NE). The filters were imaged using a LICOR Odyssey imaging system.

HeLa cells were infected with *Ct* transformed with the pBOMB4-Tet (-GFP) plasmid encoding mCherry-PBP2 or mCherry-PBP3. The fusions were induced with 10nM aTc at 17hpi. Uninduced cells were included as a control. The cells were harvested at 21hpi, and samples were processed for immunoblotting as described above. The blots were probed with rabbit polyclonal antibodies raised against peptides derived from chlamydial PBP2 or PBP3 (Ouellette SP 2012). The blots were then rinsed and incubated with 800 donkey anti-rabbit IgG secondary antibodies (LICOR, Lincoln, NE). The filters were imaged using a LICOR Odyssey imaging system.

## ACKNOWLEDGMENTS

We thank Dr. H. Caldwell (NIH/NIAID) for providing eukaryotic cell lines and Dr. I. Clarke (University of Southampton) for providing the plasmidless strain of *C. trachomatis* serovar L2. Funding for this work was supported in part by the National Institutes of Health (NIH/NIGMS) grant R35GM151971 to SPO and by the NSF grant 1817578 to JVC.

---

**Supp. Fig. S1:**
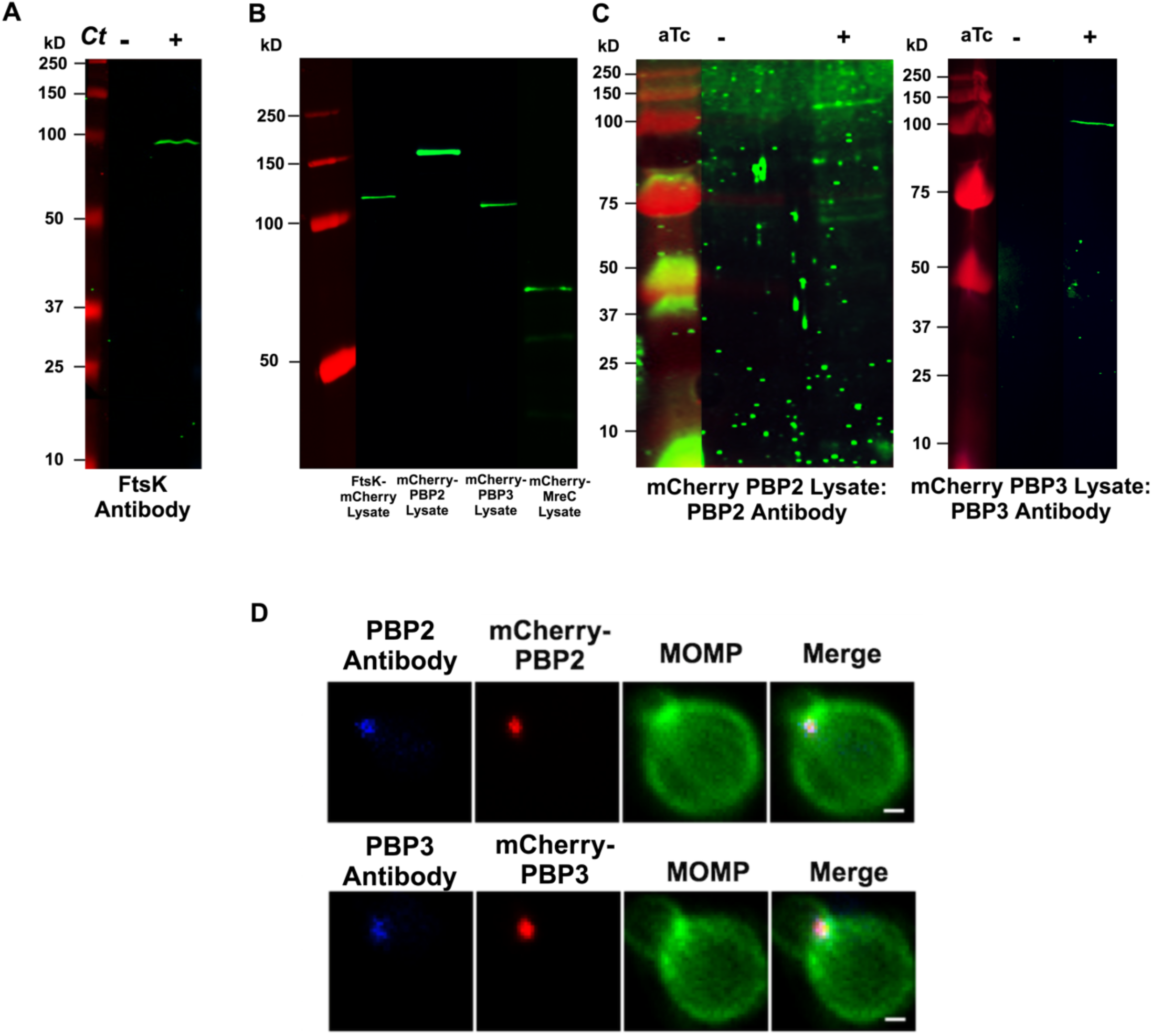
(A) Lysates were prepared from uninfected HeLa cells and HeLa cells infected with *Ct* L2. At 21hpi, lysates were prepared and characterized by immunoblotting with FtsK-specific antibodies. (B) HeLa cells were infected with *Ct* transformed with the various mCherry fusions used in this study; FtsK-mCherry (molecular mass - 114,664 Da), mCherry-PBP2 (molecular mass - 150,842 Da), mCherry-PBP3 (molecular mass - 100,150 Da), or mCherry-MreC (molecular mass - 63,906 Da). The fusions were induced with 10nM aTc at 17hpi. HeLa cells were harvested at 21hpi and lysates were prepared and characterized by immunoblotting analysis with a rabbit polyclonal mCherry antibodies. (C) HeLa cells were infected with *Ct* transformed N-terminal fusions of PBP2 or PBP3. The fusions were induced (+aTc) at17hpi. Controls were not induced (-aTc). The cells were harvested at 21hpi and lysates were prepared as described in the Methods and characterized by immunoblotting analysis with rabbit antibodies raised against peptides derived from chlamydial PBP2 or PBP3. The PBP3 antibody primarily detects a single species with the predicted molecular mass of mCherry-PBP3 in the induced sample. The PBP2 antibody primarily detects a species of ∼120kD in the induced sample, which is smaller than the predicted molecular mass of mCherry-PBP2 (∼150kD). The failure to detect full length mCherry-PBP2 may be due to the masking of the epitope recognized by the PBP2 antibody by the N-terminal mCherry tag in the full-length protein. (D) HeLa cells were infected with *Ct* transformed with mCherry-PBP2 or mCherry-PBP3. The fusions were induced by the addition of 10nM aTc to the media of the infected cells at 19hpi. Infected cells were harvested at 21hpi and lysates were prepared and stained with the PBP2 or PBP3 antibodies. The staining with the PBP2 and PBP3 antibodies completely overlaps the mCherry fluorescence from the mCherry-PBP2 and mCherry-PBP3 fusions (Bars are 3μm)

**Supp. Fig. S2:**
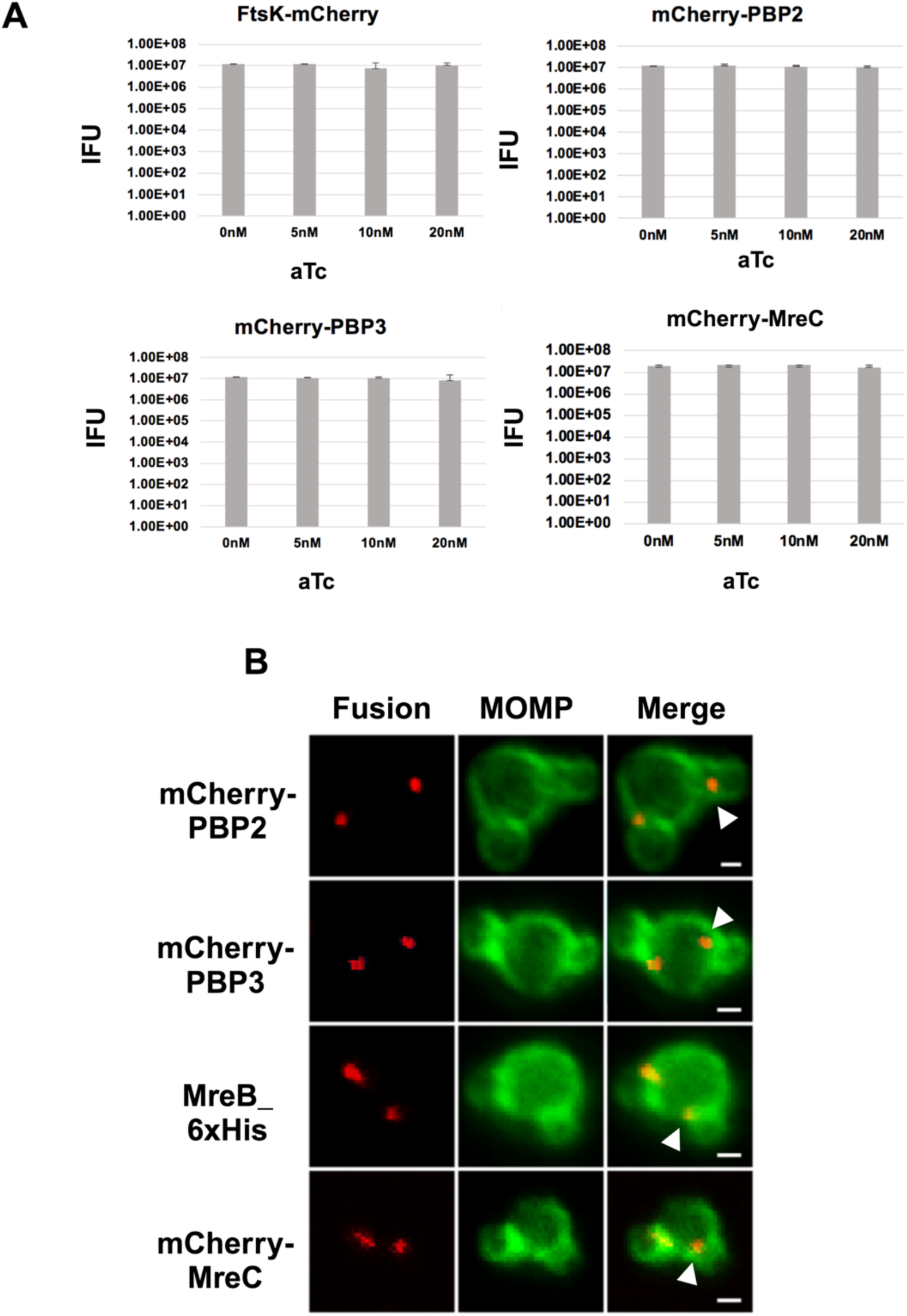
(A) HeLa cells were infected with *Ct* transformed with FtsK-mCherry, mCherry-PBP2, mCherry-PBP3, and mCherry-MreC. The fusions were uninduced or induced by the addition of varying amounts of aTc to the media of the infected cells at 8hpi. The cells were harvested at 48hpi and *Ct* were isolated. The number of infectious *Ct* in the lysates was measured by an IFU assay. Chi-squared analysis revealed that induction of the fusions did not have a statistically significant effect on the growth of *Ct* and the production of infectious EBs. (B) Each of the mCherry fusions accumulate in foci at the septum and in foci at the base (marked with arrowheads) of dividing cells with secondary buds.

**Supp. Fig. S3:**
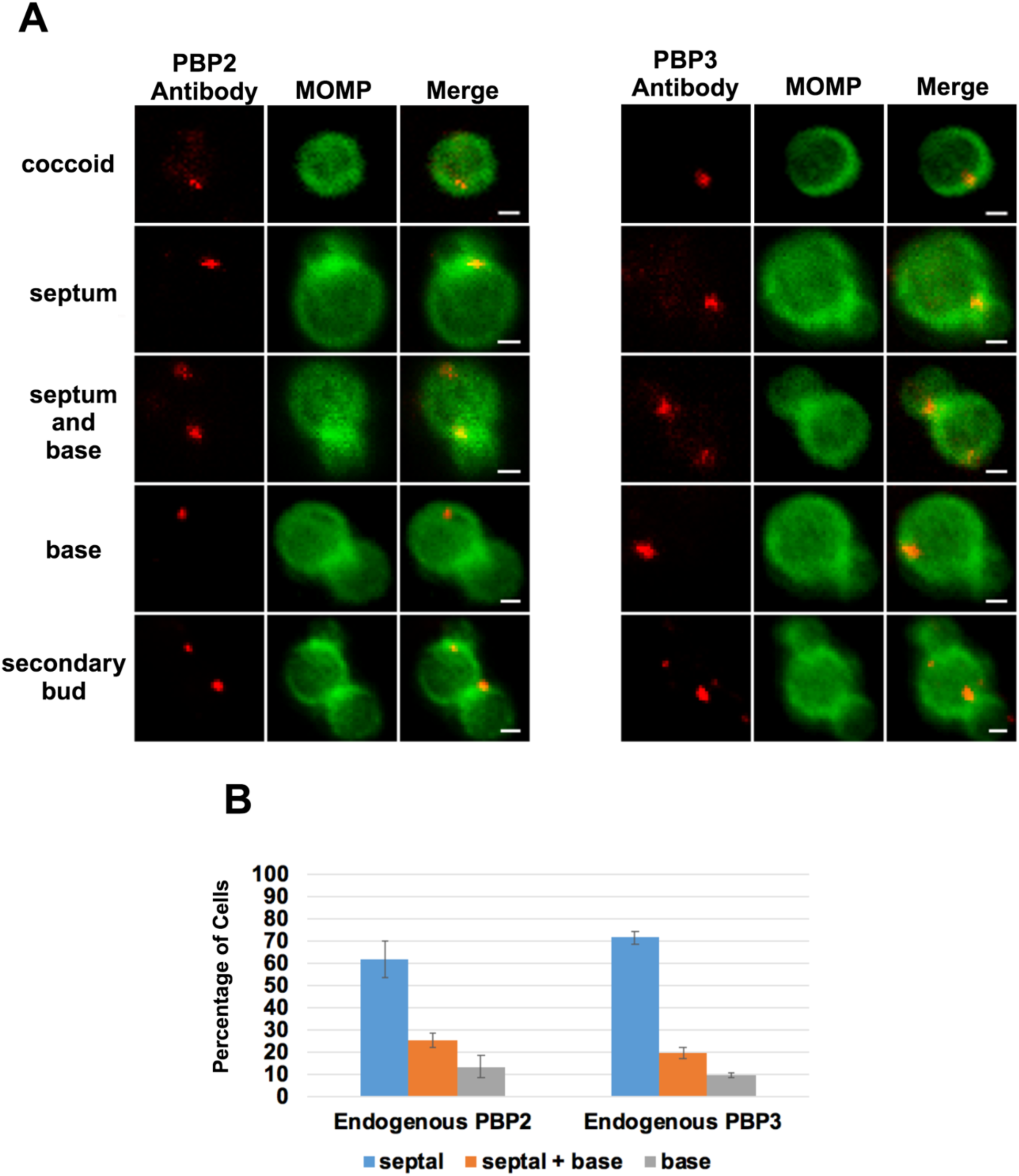
(A) Localization analyses with rabbit polyclonal antibodies that recognize endogenous PBP2 or PBP3. These analyses revealed that endogenous PBP2 and PBP3 accumulate in foci in coccoid cells, and in foci at the septum, foci at the septum and base, or in foci at the base alone in cell division intermediates in *Ct*. PBP2 and PBP3 foci are also detected at the base of secondary buds. Bars are1μm (B) Quantification revealed that the localization profiles of endogenous PBP2 and PBP3 were not statistically different than the localization profiles of the mCherry-PBP2 and mCherry-PBP3 fusions shown in Fig. 2C.

**Supp. Fig. S4:**
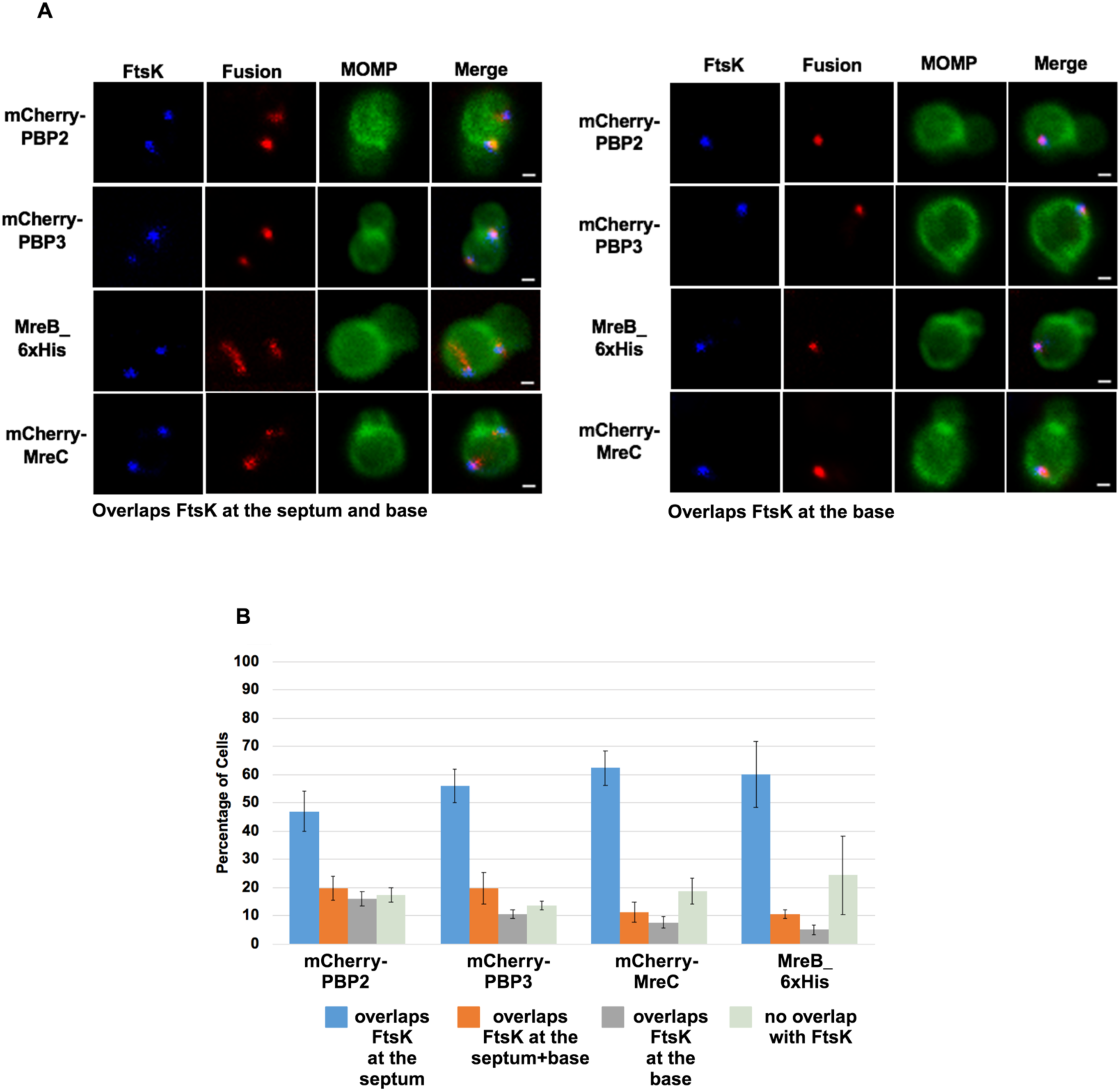
HeLa cells were infected with *Ct* transformed with PBP2, PBP3, or MreC with an N-terminal mCherry tag, or with *Ct* transformed with an MreB_6xHis fusion (Lee 2020). Each of the fusions was induced by adding 10nM aTc to the media at 17hpi. Lysates were prepared at 21hpi and the cells were fixed and stained with a MOMP antibody. The distribution of the mCherry fluorescence in dividing cells that had not initiated secondary bud formation is shown. Cells expressing the MreB_6xHis fusion were stained with rabbit anti-6xHis antibody (red) and MOMP antibodies (green). In the dividing cells shown there are (A) foci of the fusions at the septum and at the base of the mother cell or foci at the base of the mother cell only that overlap the distribution of FtsK. Bars are1μm (B) The percentage of dividing cells in which the fusions overlapped the distribution of FtsK at the septum, at the septum and at the base, and at the base alone was quantified in 100 cells. Three independent replicates were performed and the values shown are the average of the 3 replicates.

**Supp. Fig. S5:**
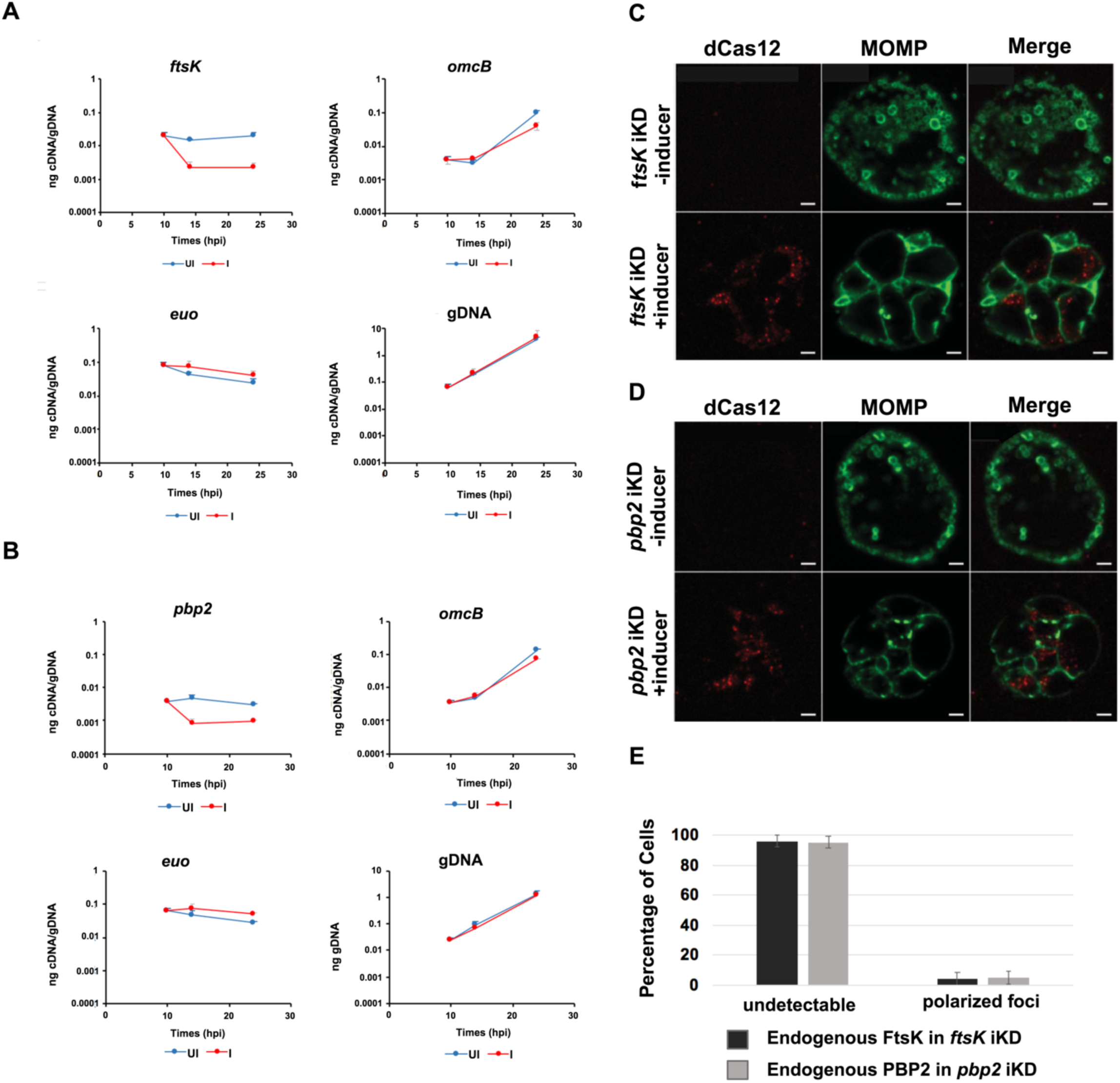
HeLa cells were infected with *Ct* transformed with the pBOMBL12CRia plasmid that constitutively expresses *ftsK* or *pbp2*-targeting crRNAs. dCas12 expression was induced by the addition of 5nM aTc to the media of infected cells at 8hpi. Control cells were not induced. Nucleic acids were isolated from induced cells and from uninduced controls at various times post-infection, and RT-qPCR was used to measure *ftsK* or *pbp2* transcript levels. (A) The induction of dCas12 resulted in ∼10-fold reduction in *ftsK* transcript levels in cells expressing the *ftsK*-targeting crRNA, (B) and ∼8-fold reduction in *pbp2* transcript levels in cells expressing the *pbp2*-targeting crRNA, while these crRNAs had minimal or no effect on chlamydial *euo* and *omcB* transcript levels. HeLa cells were infected with *Ct* transformed pBOMBL12CRia plasmid that constitutively expresses a (C) *ftsK* or (D) *pbp2*-targeting crRNA. dCas12 expression was induced by the addition of 5nM aTc to the media of infected cells at 8hpi. Control cells were not induced. The infected cells were fixed at 24hpi and stained with MOMP and Cas12 antibodies. *Ct* morphology was normal and dCas12 was undetectable in the inclusions of uninduced control cells. Foci of dCas12 were observed in induced cells, and *Ct* in the inclusion exhibited an enlarged aberrant morphology. Bars in C and D are 2μm. (E) HeLa cells were infected with *Ct* transformed with the pBOMBL12CRia plasmid that constitutively expresses a *ftsK* or *pbp2*-targeting crRNA. dCas12 was induced at 17hpi by the addition of 10nM aTc to the media. Control cells were not induced. The cells were harvested at 21hpi, and *Ct* were prepared and stained with antibodies that recognize that endogenous FtsK or PBP2. Quantification shows that polarized foci of FtsK and PBP2 were almost undetectable when *ftsK* or *pbp2* were transiently knocked down.

**Supp. Fig. S6.**
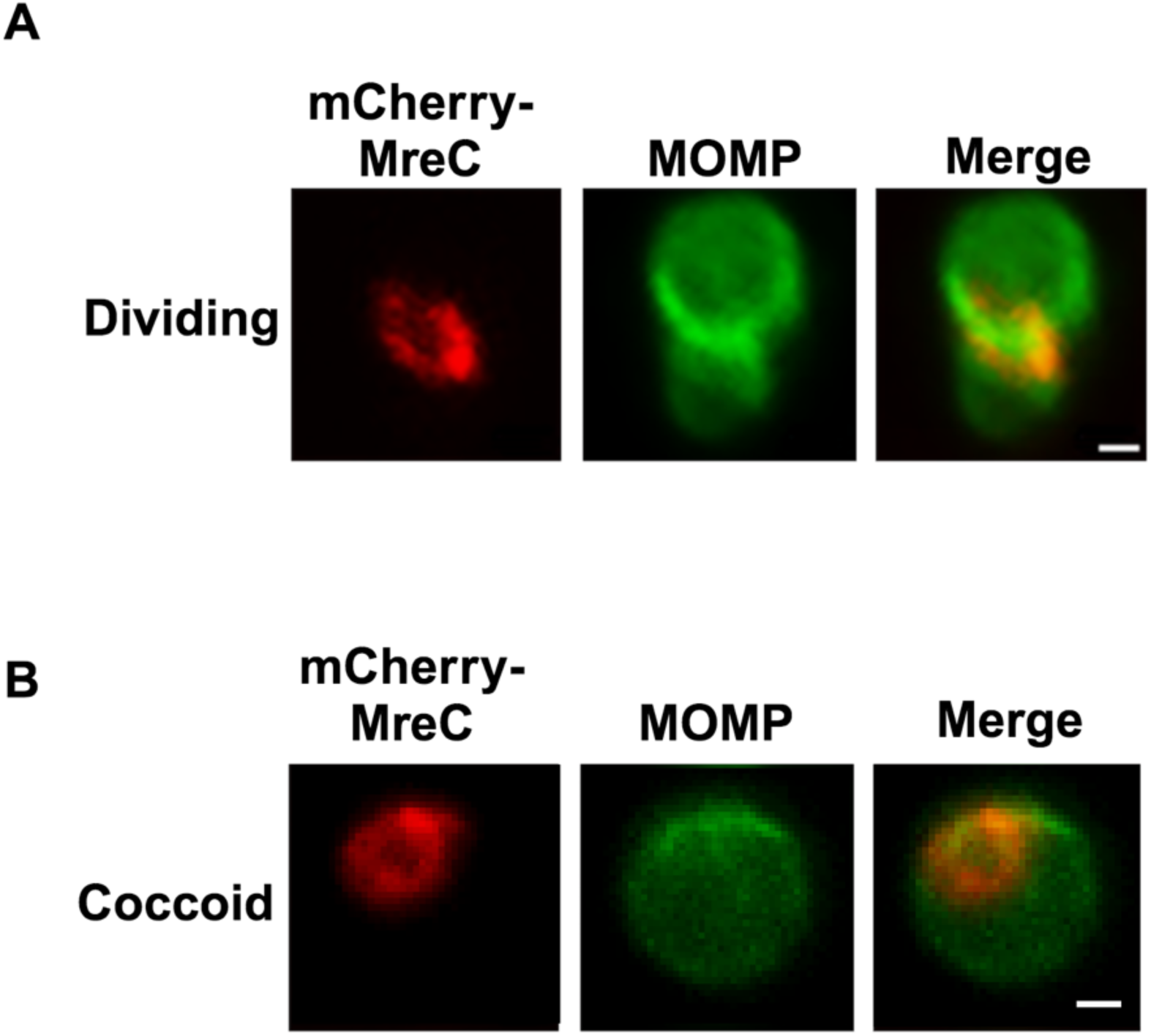
MreC rings in (A) dividing *Ct* and in (B) coccoid *Ct*. Bars are1μm.

**Supp. Table S1.**
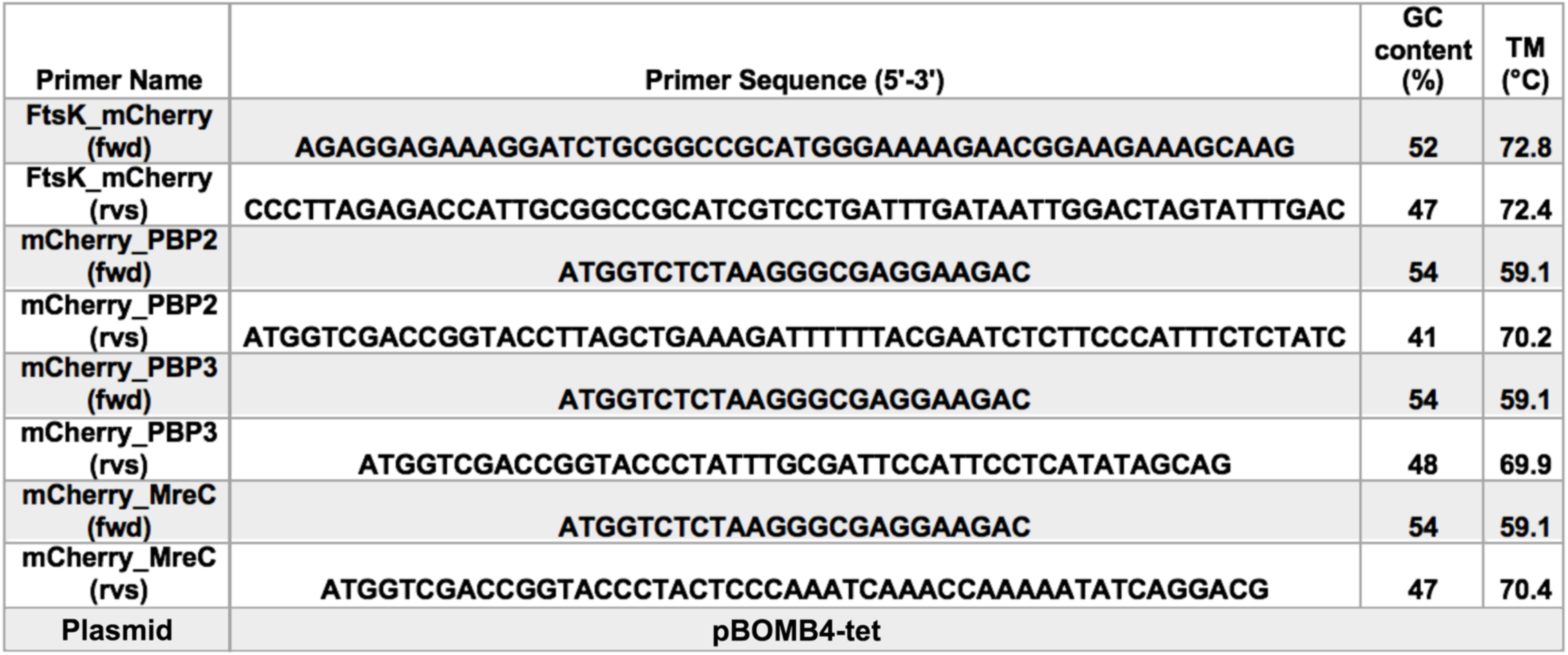
List of primers and plasmids used for cloning mCherry fusions of FtsK, PBP2, PBP3, MreB and MreC.

**Supp. Table S2:**
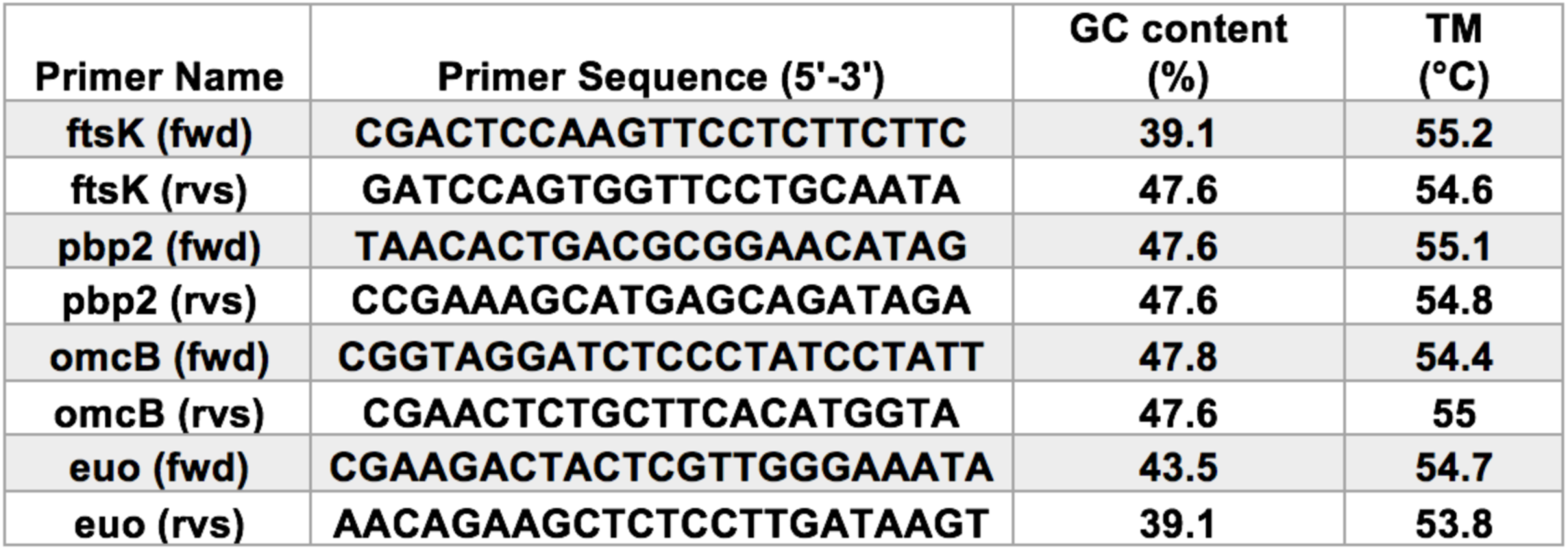
List of primers used for RT-qPCR.

